# Light-triggered and phosphorylation-dependent 14-3-3 association with NONPHOTOTROPIC HYPOCOTYL 3 is required for hypocotyl phototropism

**DOI:** 10.1101/2021.04.09.439179

**Authors:** Lea Reuter, Tanja Schmidt, Prabha Manishankar, Christian Throm, Jutta Keicher, Andrea Bock, Claudia Oecking

**Affiliations:** Center for Plant Molecular Biology (ZMBP), Plant Physiology, University of Tübingen, Germany

## Abstract

<u>N</u>ON-PHOTOTROPIC HYPOCOTYL 3 (NPH3) is a key component of the phototropic response, acting downstream of the primary photoreceptor phototropin and upstream of auxin redistribution. Despite the obvious physiological significance of the blue light-induced differential growth process, the molecular mode of NPH3 action is poorly understood. Light-triggered dephosphorylation of NPH3, however, is thought to constitute a major signaling event. Here, we show that NPH3 directly binds to polyacidic phospholipids via a polybasic motif in its C-terminal domain, allowing for plasma membrane association in darkness. We further demonstrate that blue light induces phosphorylation of a C-terminal 14-3-3 binding motif in NPH3. Subsequent binding of 14-3-3 to the phosphorylated NPH3 in turn is required for light-triggered release of NPH3 from the plasma membrane. In the cytosol, NPH3 undergoes a dynamic transition from a dilute to a condensed state. Intriguingly, the dephosphorylated state of the 14-3-3 binding site as well as NPH3 plasma membrane association are recoverable in darkness. Given that NPH3 variants constitutively localizing either to the plasma membrane or to cytosolic condensates are non-functional, the phototropin-triggered and 14-3-3 mediated dynamic change in the subcellular localization of NPH3 seems to be crucial for its function. Taken together, our data demonstrate a fundamental role for 14-3-3 members in regulating NPH3 localization and auxin-dependent phototropic responses.

## Introduction

Developmental plasticity of plants is impressively demonstrated by the phototropic response, through which plants align their growth with incoming blue light (BL) [1]. Shoots typically grow towards the light by generating a lateral gradient of the growth promoting phytohormone auxin. Here, the hormone concentration is higher on the shaded side as compared with the lit side, resulting in differential growth. It is well established that the phototropins phot1 and phot2 function as primary photoreceptors controlling phototropism in *Arabidopsis*: [2-4]. Phototropins are plasma membrane (PM)-associated, light-activated protein kinases and indeed, blue-light (BL) induced autophosphorylation turned out to be a primary and essential step for the asymmetric growth response [5]. Interestingly, members of the 14-3-3 family have been identified as phot1 interactors. 14-3-3 proteins are known to interact with a multitude of polypeptides in a phosphorylation-dependent manner, thereby regulating distinct cellular processes [6]. As yet, however, a functional role of phot1/14-3-3 association could not be proven [5, 7]. Furthermore, evidence for trans-phosphorylation activity of phototropins is surprisingly limited.

The polar localization of PIN auxin efflux carriers within the PM made them likely candidates promoting formation of the auxin gradient that precedes phototropic growth [8]. Indeed, a mutant lacking the three major PINs expressed in aerial plant parts (PIN3, PIN4, PIN7) is severely compromised in phototropism [9]. Notably, unilateral illumination polarizes PIN3 specifically to the inner lateral side of hypocotyl endodermis cells, aligning PIN3 polarity with the light direction and presumably redirecting auxin flow towards the shaded side [10]. Moreover, the activity of PINs is positively regulated by two protein kinase families from the AGCVIII class, namely PINOID and D6 PROTEIN KINASES [11]. Though phototropins belong to the same kinase class, direct PIN phosphorylation could not be shown [10]. Taken together, signaling events that couple photoreceptor activation to changes in PIN polarization and consequently auxin relocation remain mainly elusive.

In this regard, the PM-associated NON-PHOTOTROPIC HYPOCOTYL 3 (NPH3) might represent a promising component of early phototropism signaling events. It acts downstream of the photoreceptors and appears to be instrumental for auxin redistribution [3, 4, 12, 13]. NPH3 possesses – in addition to the central NPH3 domain – two putative protein-protein interaction domains, a C-terminal coiled-coil (CC) domain and a N-terminal BTB/POZ (<u>b</u>road-complex, tramtrack, <u>b</u>ric a brac/<u>Po</u>x virus and <u>z</u>inc finger) domain [1, 14]. Indeed, NPH3 physically interacts not only with the photoreceptor phot1 but also with further early signaling elements, such as ROOT PHOTOTROPISM (RPT2) [15] - another member of the plant-specific NPH3/RPT2-like family (NRL) - and defined members of the PHYTOCHROME KINASE (PKS) family [16, 17]. Interestingly, NPH3 exists in a phosphorylated form in dark-grown seedlings and becomes rapidly dephosphorylated upon phot1 activation [18, 19]. Later on, the alteration in phosphorylation status was shown to correlate closely with light-driven changes in the subcellular localization of NPH3 which detaches from the PM upon irradiation, forming aggregated particles in the cytosol [20]. As found for light-triggered dephosphorylation [18], formation of the NPH3 particles is reversible upon darkness or prolonged irradiation [20]. One factor required for the recovery of phosphorylated NPH3 at the PM over periods of prolonged irradiation is its interaction partner RPT2 [20]. Altogether, this has led to the current model that the phosphorylation status of NPH3 determines its subcellular localization and function: phosphorylation of NPH3 promotes its action in mediating phototropic signaling from the PM, whereas NPH3 dephosphorylation reduces it by internalizing NPH3 into aggregates [4, 7, 12, 20]. As yet, however, the functional significance of NPH3 (de)phosphorylation remains poorly understood [19, 21].

Here, we identified members of the 14-3-3 family as novel interactors and major regulators of NPH3. Our analyses revealed that BL induces phosphorylation of the antepenultimate NPH3 residue which in turn enables 14-3-3 association. Complex formation interferes with the ability of NPH3 to bind to polyacidic phospholipids, resulting in its displacement from the PM. Accumulation of NPH3 in the cytosol causes formation of membrane-less condensates. Intriguingly, both PM association and 14-3-3 triggered PM dissociation are required for NPH3 function. Taking the reversibility of the light-induced processes into account, the phototropin-regulated and 14-3-3-mediated dynamic change in the subcellular localization of NPH3 seem to be crucial for its proper function in the phototropic response.

## Results

### 14-3-3 proteins interact with NPH3 via a C-terminal binding motif in a BL-dependent manner

A yeast two hybrid screen performed in our lab (see [22]) identified NPH3 as putative interactor of several Arabidopsis 14-3-3 isoforms, among those epsilon and omega (Fig. 1A). Complex formation of NPH3 and 14-3-3 was confirmed *in planta* by co-immunoprecipitation (CoIP) of fluorophore-tagged proteins transiently co-expressed in *N. benthamiana* leaves (Fig. 1B). To elucidate the impact of light on 14-3-3/NPH3 complex assembly, transgenic Arabidopsis lines expressing 14-3-3 epsilon:GFP under control of the native promoter [23] and, as control, UBQ10::GFP were employed. Three-days old etiolated seedlings were either maintained in complete darkness or irradiated with BL (1μmol m-^2^sec^-1^) for 30 minutes.

**Fig. 1:**
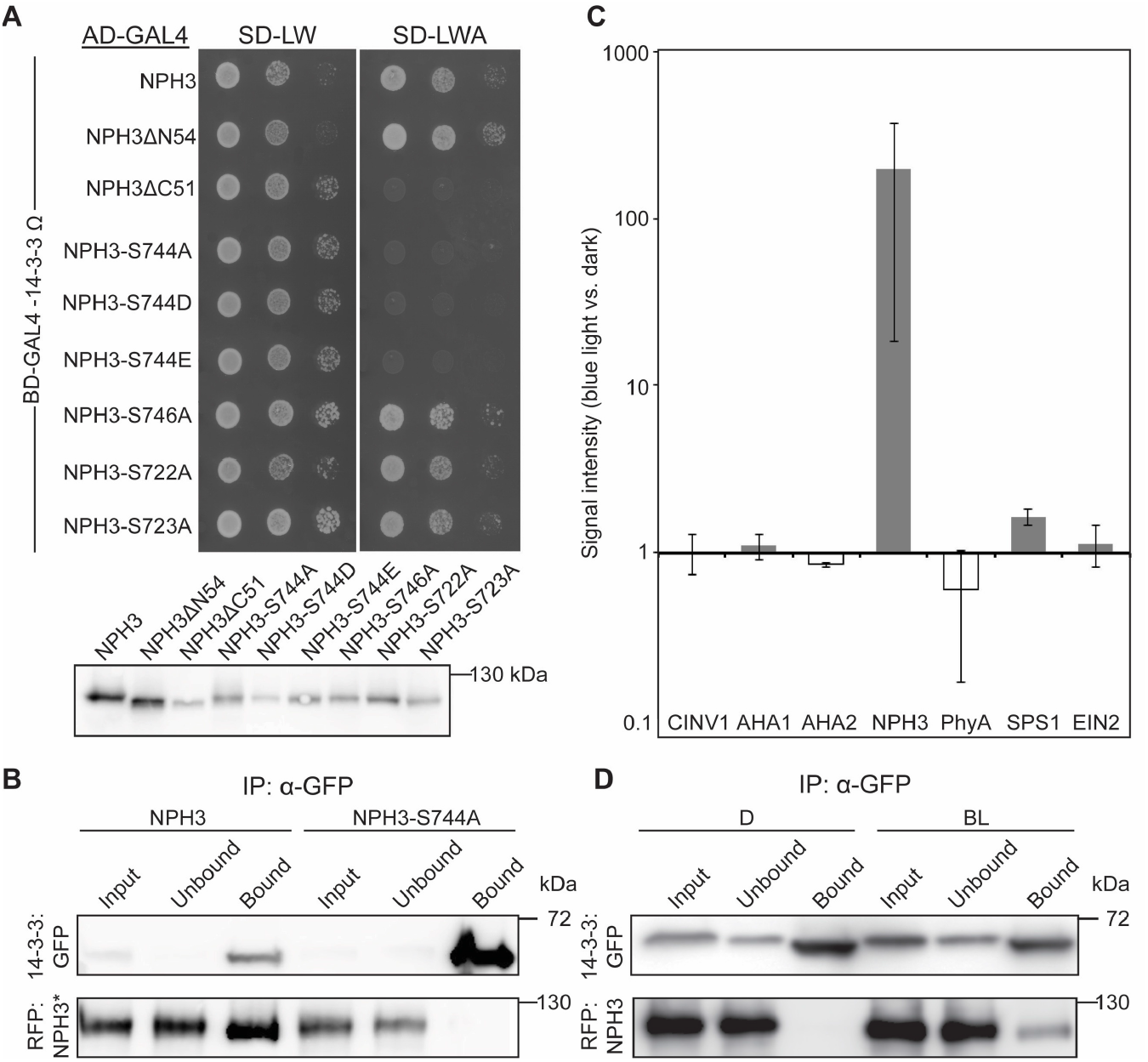
Interaction of NPH3 and 14-3-3 proteins is triggered by blue light irradiation and abolished by mutation of the antepenultimate NPH3 residue. **(A)** Yeast two-hybrid interaction analysis of the Arabidopsis 14-3-3 isoform omega with NPH3 wild type and mutant variants (upper panel). Expression of the diverse NPH3 fusion proteins in yeast was confirmed by anti-HA-immunodetection (lower panel). AD, activating domain; BD, binding domain. **(B)** *In vivo* interaction of mCherry:NPH3 variants and 14-3-3 omega:mEGFP in transiently transformed *N. benthamiana* leaves. Expression of transgenes was driven by the 35S promoter. Freshly transformed tobacco plants were kept under constant light for 42 h. The crude extract was immunoprecipitated using GFP beads and separated on 11% SDS-PAGE gels, followed by immunoblotting with anti-GFP and anti-RFP antibodies, respectively. **(C)** Arabidopsis 14-3-3 epsilon interactors were identified by mass spectrometry analysis of anti-GFP immunoprecipitations (two biological replicates) from etiolated seedlings expressing 14-3-3 epsilon:GFP either maintained in darkness or irradiated with blue light (1 μmol m^-2^ sec^-1^) for 30 min. Protein intensities of well-known 14-3-3 client proteins were normalized to relative abundance of the bait protein (Table S1). Fold changes in relative abundance (mean ± SD, logarithmic scale) of blue light treatment versus darkness are given. AHA1, AHA2, Arabidopsis H^+^-ATPase; CINV1, cytosolic invertase 1; EIN2, ethylene insensitive 2; PhyA, phytochrome A; SPS1, sucrose phosphate synthase 1. **(D)** Light dependence of the *in vivo* interaction of mCherry:NPH3 and 14-3-3 omega:mEGFP in transiently transformed *N. benthamiana* leaves. Expression of transgenes was driven by the 35S promoter. Freshly transformed tobacco plants were kept under constant light for 24 h and subsequently transferred to darkness for 17h with (BL) or without (D) blue light treatment (5 μmol m^-2^ sec^-1^) for the last 40 minutes. The crude extract was immunoprecipitated using GFP beads and separated on 11% SDS-PAGE gels, followed by immunoblotting with anti-GFP and anti-RFP antibodies, respectively.

Potential targets of 14-3-3 epsilon:GFP were identified by stringent CoIP-experiments coupled with mass spectrometry (MS)-based protein identification. As expected, several known 14-3-3 clients [23] were detected by MS, and remarkably, NPH3 emerged as a major 14-3-3 interactor (Table S1). Binding capability of characterized 14-3-3 targets, such as the H^+^-ATPase (AHA1) and cytosolic invertase 1 (CINV1), was not modified by BL treatment. By contrast, NPH3 turned out to be a BL-dependent 14-3-3 interactor *in planta* (Fig. 1C, Table S1). CoIP of fluorophore-tagged proteins transiently co-expressed in *N. benthamiana* leaves confirmed that physical association of NPH3 and 14-3-3 is not detectable in darkness while BL irradiation triggers complex formation (Fig. 1D). Assuming 14-3-3 association to depend on phosphorylation of the target protein, this observation is in apparent contrast to the light-induced dephosphorylation of NPH3 [18].

The specific phosphorylatable 14-3-3 binding sequences of numerous target proteins are most flexible and disordered [24]. Since both the N- and C-terminal domain of NPH3 are predicted to be intrinsically disordered (Fig. S1, [25]), the corresponding truncated versions were analyzed by yeast two hybrid assays. While NPH3 lacking the N-terminal 54 residues (NPH3ΔN54, still comprising the BTB domain) was capable of 14-3-3 binding, deletion of the C-terminal 51 residues (NPH3ΔC51, still comprising the CC domain) abolished 14-3-3 association, suggesting that the 14-3-3 binding site localizes downstream of the CC domain (Fig.1A). We therefore exchanged amino acid residues, phosphorylation of which has recently been demonstrated *in planta* (S722, S723, S744, S746, [26, 27]), for a non-phosphorylatable alanine. Strikingly, 14-3-3 binding was not affected in all but one NPH3 mutant: replacement of S744 – the antepenultimate residue of NPH3 – prevented 14-3-3 association both in yeast (Fig. 1A) and *in planta* (Fig. 1B), suggesting a phosphorylation-dependent C-terminal 14-3-3 binding motif (pS/pTX_1-2_-COOH) [28] in NPH3. Phosphomimic variants (NPH3-S744D/S744E), however, do not allow for 14-3-3 binding (Fig. 1A), a characteristic of almost all 14-3-3 clients [29].

### 14-3-3 association is required for NPH3 function and its BL-induced PM dissociation

To address the issue of functional significance of 14-3-3 association *in vivo*, GFP-tagged NPH3 variants were expressed in a T-DNA induced *loss of function* allele of *NPH3, nph3-7* [30]. GFP:NPH3 was fully functional in restoring the severe impairment of hypocotyl phototropism in *nph3-7*, regardless of whether expression was driven by the native or the 35S CaMV promoter (Fig. 2A, Fig. S2A), thus confirming previous data [7, 20]. By contrast, phototropic hypocotyl bending was still significantly reduced when NPH3 incapable of 14-3-3 association (GFP:NPH3-S744A) was expressed (Fig. 2A, Fig. S2A), indicating that BL-induced interaction with 14-3-3 is required for proper NPH3 function.

**Fig. 2:**
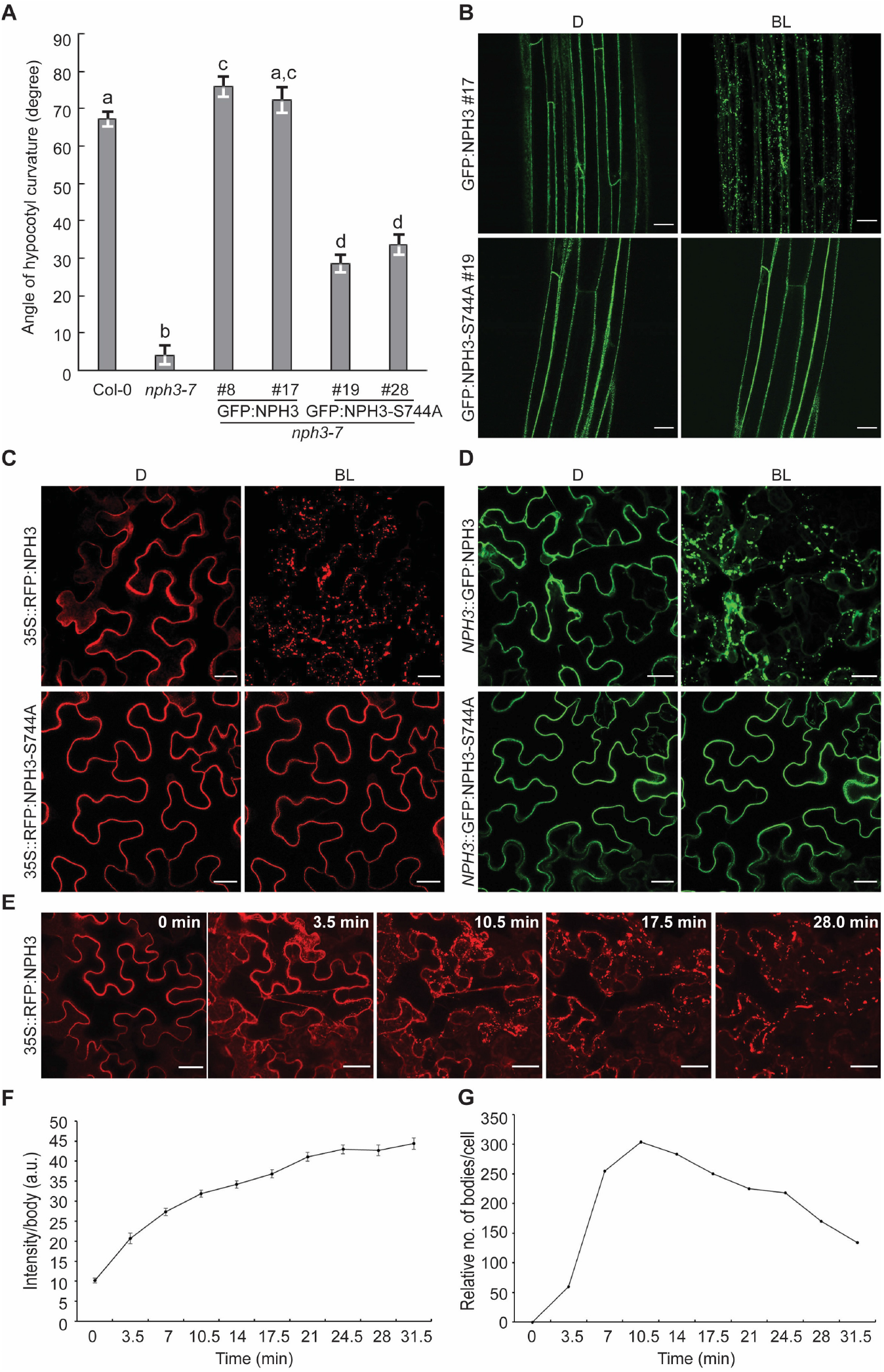
14-3-3 binding is required for proper NPH3 function in phototropic hypocotyl bending and its light-triggered detachment from the plasma membrane. **(A)** Quantification of the hypocotyl phototropism response (mean ± SEM) in 3-days old etiolated seedlings exposed for 12h to unilateral blue light (1 μmol m^-2^ sec^-1^) (n>30 seedlings per experiment, one representative experiment of two replicates is shown). Expression of transgenes in *nph3-7* was driven by the 35S promoter. Student’s t-test, different letters mark statistically significant differences (P<0.05), same letters mark statistically non-significant differences. **(B, C, D**) Representative confocal microscopy images of hypocotyl cells from transgenic etiolated Arabidopsis *nph3-7* seedlings **(B)** or of leaf epidermal cells from transiently transformed *N. benthamiana* (Z-stack projections of BL-treated NPH3 are shown) **(C, D)**. The plants were either kept in darkness (D) or treated with blue light (BL) (*N. benthamiana*: approx. 11 min and *nph3-7*: approx. 6 min by means of the GFP-laser). Expression of transgenes was driven by the 35S promoter **(B, C)** or the native *NPH3* promoter **(D)**. Scale bars, 25 μm. **(E, F, G)** Single-cell time-lapse imaging of RFP:NPH3 condensation induced by GFP-laser treatment. The image of time point 0 image was taken in the absence of the GFP-laser. Z-stack projections from selected time points **(E)**, fluorescence intensity per body (mean ± SEM) **(F)** and number of bodies **(G)** are shown. Scale bars, 25 μm.

Though NPH3 is hydrophilic in nature, both GFP:NPH3 and GFP:NPH3-S744A localized to the cell periphery in the hypocotyl of etiolated transgenic seedlings (Fig. 2B, Fig. S2B), suggesting PM association as described previously for NPH3 [1, 7, 20]. Within minutes, however, the BL laser used to excite GFP (488 nm, activates phototropins) induced detachment of NPH3 from the PM into discrete bodies/particle-like structures in the cytoplasm (Video S1). This BL-induced shift in subcellular localization is mediated by phot1 activity [20] and again, could be observed independent of whether expression of NPH3 was under control of the endogenous (Fig. S2B; [7]) or the 35S promoter (Fig. 2B, [20]). By contrast, GFP:NPH3-S744A remained mainly PM-associated upon irradiation (Fig. 2B, Video S2, Fig. S2B,C). Mutation of the 14-3-3 binding site does thus not affect PM association of NPH3 in darkness but prevents BL-triggered PM dissociation, suggesting that light-induced association of 14-3-3 proteins is required to internalize NPH3 from the PM into cytosolic particles. We confirmed our findings in transiently transformed *N. benthamiana* leaves (Fig. 2C, D; Videos S3, S4). Here, primarily RFP-tagged proteins were employed since excitation of RFP (558 nm) – unlike GFP (488 nm) – does not activate phototropins. This enabled us to conditionally activate phot1 by means of the GFP laser. It became evident that NPH3 – instead of being directly internalized into discrete bodies - initially detaches from the PM and moves along cytoplasmic strands comparable to soluble polypeptides (Video S3). Body formation in the cytosol is initiated after a lag time of approximately 4 to 5 minutes. Generation of particle-like structures might thus depend on soluble NPH3 exceeding a critical concentration in the cytosol. Upon co-expression of GFP-tagged 14-3-3s, colocalization with NPH3 was observed in discrete bodies (Fig. S2D).

### PM association of NPH3 is phospholipid-dependent and requires its C-terminal domain

According to the data presented so far, light-induced 14-3-3 association interferes with PM anchoring of NPH3 (Fig. 2B-D). Association of the hydrophilic NPH3 to the PM is known since its discovery in 1999 [1]. As yet, however, the molecular determinants of membrane recruitment of NPH3 remain elusive. Proteins can be anchored to membranes by diverse mechanisms such as hydrophobic interactions, protein-protein and protein-lipid interactions. In eukaryotes, the PM is the most electronegative compartment of the cell. In plants, the phosphoinositide phosphatidylinositol-4-phosphate (PI4P), phosphatidic acid (PA) and phosphatidylserine (PS) are separately required to generate the strong and unique electrostatic signature of the PM [31-34]. To examine the importance of electronegativity for NPH3 PM association, we made use of a genetic system that depletes the polyacidic PI4P at the PM by PM-anchoring of the catalytic domain of the yeast SAC1 PI4P phosphatase [33, 35]. Transient co-expression of NPH3 together with SAC1, but not the catalytically inactive version SAC1_DEAD_, displaced NPH3 from the PM into discrete cytosolic bodies in darkness, thus reflecting the status typically induced by BL treatment (Fig. 3A). In lipid overlay assays, NPH3 bound to several phospholipids characterized by polyacidic headgroups, namely PA as well as the phosphoinositides PI3P, PI4P, PI5P, PI(3,4)P_2_, PI(3,5)P_2_, PI(4,5)P_2_ and PI(3,4,5)P_3_. NPH3 did neither bind to phospholipids with monoacidic headgroups such as phosphatidylinositol (PI) or PS, nor to phospholipids with neutral headgroups, namely phosphatidylcholine (PC) and phosphatidylethanolamine (PE) (Fig. 3B). NPH3ΔC51, however, was unable while the bacterially expressed C-terminal 51 residues of NPH3 (NPH3-C51) turned out to be sufficient to bind to polyacidic phospholipids (Fig. 3B). Furthermore, NPH3-C51 bound to large unilamellar liposomes containing the polyacidic phospholipids PI4P or PA, but not to liposomes composed of only neutral phospholipids such as PC and PE (Fig. 3C). Obviously, the C-terminal 51 residues of NPH3 enable electrostatic association with membrane bilayers irrespective of posttranslational protein modifications or association with other proteins. Accordingly, significantly impaired PM association of NPH3ΔC51 was expected *in vivo*. Indeed, transient expression in *N. benthamiana* revealed loss of PM recruitment of GFP:NPH3ΔC51 in the dark, as evident by the presence of discrete bodies in the cytosol (Fig. 3D, Video S5, Fig. S3). This resembles the scenario observed upon co-expression of NPH3 and SAC1 as well as upon transient expression of NPH3ΔC65:GFP in guard cells of *Vicia faba* [36]. By contrast, deletion of the N-terminal domain (NPH3ΔN54) did not affect PM association of NPH3 in darkness (Fig. 3D, Video S6, Fig. S3).

**Fig. 3:**
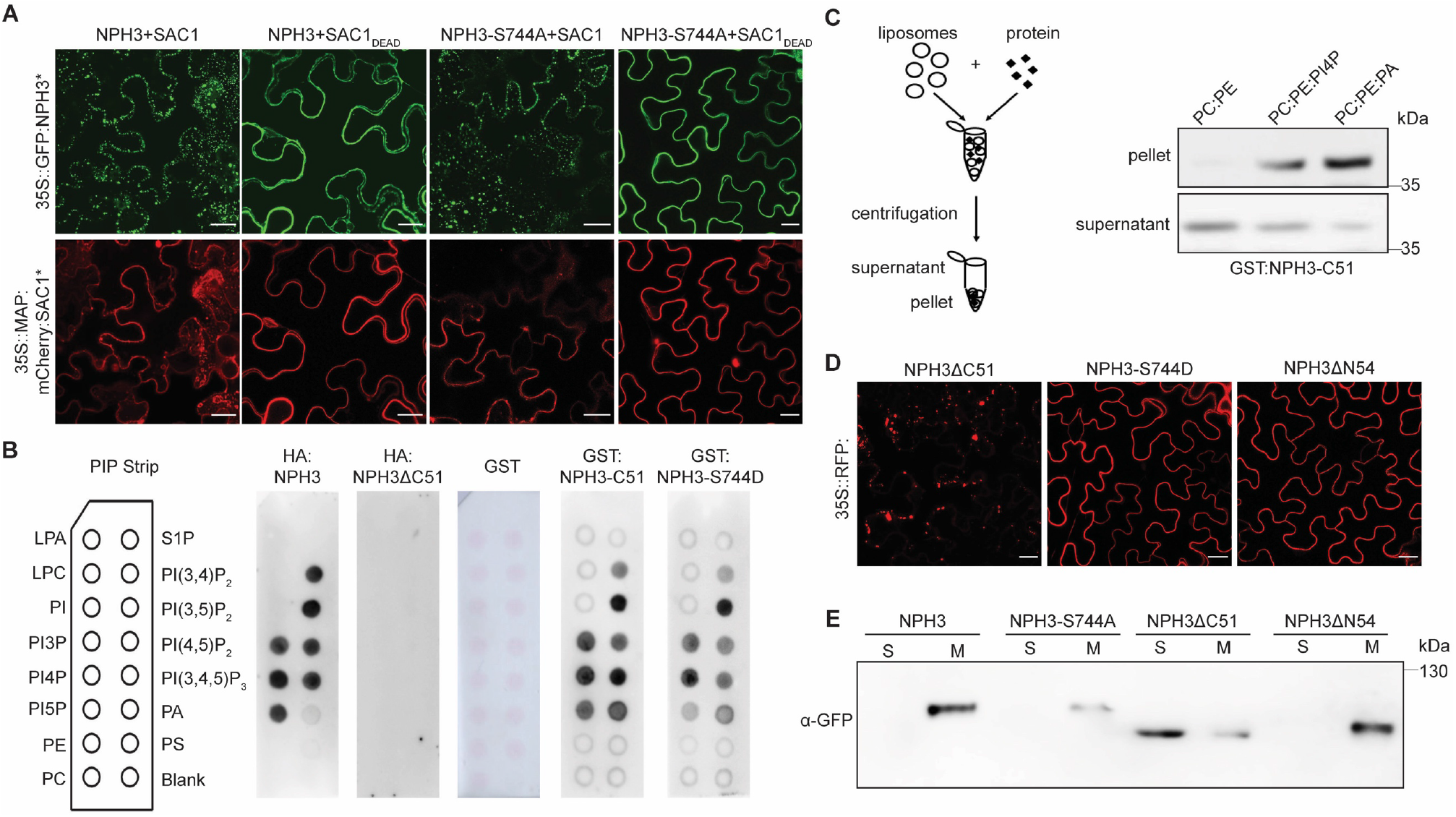
NPH3 binds to polyacidic phospholipids via its C-terminal domain. **(A), (D)** Representative confocal microscopy images of leaf epidermal cells from transiently transformed *N. benthamiana* adapted to darkness (Z-stack projections of NPH3ΔC51 **(D)** as well as NPH3 variants (NPH3*) co-expressed with SAC1 variants (SAC1*) **(A)** are shown). Scale bars, 25 μm. **(B)** Lipid overlay assay performed with either *in vitro* transcribed and translated HA:NPH3 and HA:NPH3ΔC51 or purified GST and GST:NPH3-C51 variants. Immunodetection was performed by using anti-HA or anti-GST antibodies, respectively. See main text for abbreviations. **(C)** Liposome binding assay using large unilamellar liposomes containing the neutral phospholipids PE and PC mixed with either the polyacidic PI4P or PA as specified. Anti-GST immunoblot of GST:NPH3-C51 is shown. **(E)** Representative immunoblots with anti-GFP after subcellular fractionation of protein extracts prepared from *N. benthamiana* leaves transiently expressing 35S::GFP:NPH3 variants and adapted to darkness. Proteins in each fraction (7.5 μg) were separated on 7.5% SDS-PAGE gels. Note that the total amount of soluble proteins (S) is approximately 15 times higher as compared to the total amount of microsomal proteins (M) after 100,000 g centrifugation.

Altogether, the C-terminal domain plays a dual role in determining the subcellular localization of NPH3: it mediates phospholipid-dependent PM association and allows for PM dissociation as a result of 14-3-3 association. Nonetheless, the suspected phosphorylation of S744 might *per se* decrease the interaction of NPH3 with polyacidic phospholipids, hence triggering PM dissociation. Yet, the appropriate phosphomimic version of NPH3 (NPH3-S744D) was neither impaired in phospholipid-interaction *in vitro* (Fig. 3B) nor PM recruitment *in vivo* (Fig. 3D). Consequently, association of 14-3-3 to the antepenultimate, presumably phosphorylated residue S744 seems to be crucial for the release of the NPH3 C-terminal domain from the PM.

### An amphipathic helix is essential for phospholipid binding and PM association of NPH3 *in vivo*

Several members of the AGCVIII kinase family - though not phot1-contain a basic and hydrophobic (BH) motif in the middle domain of the kinase. This polybasic motif interacts directly with phospholipids and is required for membrane binding [37]. When we applied the BH score prediction to NPH3 (window size 11 as recommended for the detection of motifs closer to the termini, [38]), two positively charged motifs in the C-terminal domain with a BH score above the critical threshold value of 0.6 were identified: (i) a R-rich motif (R736-R742) adjacent to the 14-3-3 binding site (S744) and (ii) a K-rich motif further upstream (W700-M713) (Fig. 4A, Fig. S4A). The latter is predicted to form an amphipathic helix, organized with clearly distinct positively charged and hydrophobic faces. The hydrophobic moment – a measure of the amphiphilicity - was calculated to be 0.58 (Fig. S4B), similar to the PM anchor of Remorin [39]. In order to test the requirement of the two motifs for membrane association, all five basic amino acids within the R-rich motif were replaced by alanine (NPH3-5KR/A). Furthermore, both hydrophobicity and positive charge of the amphipathic helix were decreased by exchange of four hydrophobic residues (NPH3-4WLM/A) and of four lysine residues (NPH3-4K/A), respectively (Fig. 4A; Fig. S4A). The ability of any of the three NPH3 replacement variants to bind to polyacidic phospholipids *in vitro* was significantly impaired (Fig. 4B, C). Against expectations, the GFP:NPH3-5KR/A mutant remained PM-associated in the dark when transiently expressed in *N. benthamiana* (Fig. 4D). To verify that the terminal R-rich motif directly upstream of the 14-3-3 binding site is dispensable for PM recruitment *in vivo*, NPH3 was truncated by the C-terminal 28 residues (NPH3ΔC28). Indeed, PM anchoring in the dark was unaffected (Fig. 4D). By contrast, modification of either the amphiphilicity or the hydrophobicity of the amphipathic helix gave rise to cytosolic particle-like structures in darkness (Fig. 4D, Fig. S4C). Though these particles differ in shape and size, strict co-localization of the corresponding NPH3 variants was observed upon co-expression (Fig. S4D). Taken together, these experiments reveal the necessity of the amphipathic helix for PM anchoring *in vivo*. Obviously, PM-association of NPH3 is not limited to phospholipid binding but also involves hydrophobic interactions. Thus, one attractive hypothesis is that the positively charged residues interact electrostatically with polyacidic phospholipids of the PM followed by partial membrane penetration. By this means, interactions with both the polar headgroups and the hydrocarbon region of the bilayer would be established in darkness, causing anchor properties of NPH3 similar to intrinsic proteins.

**Fig. 4:**
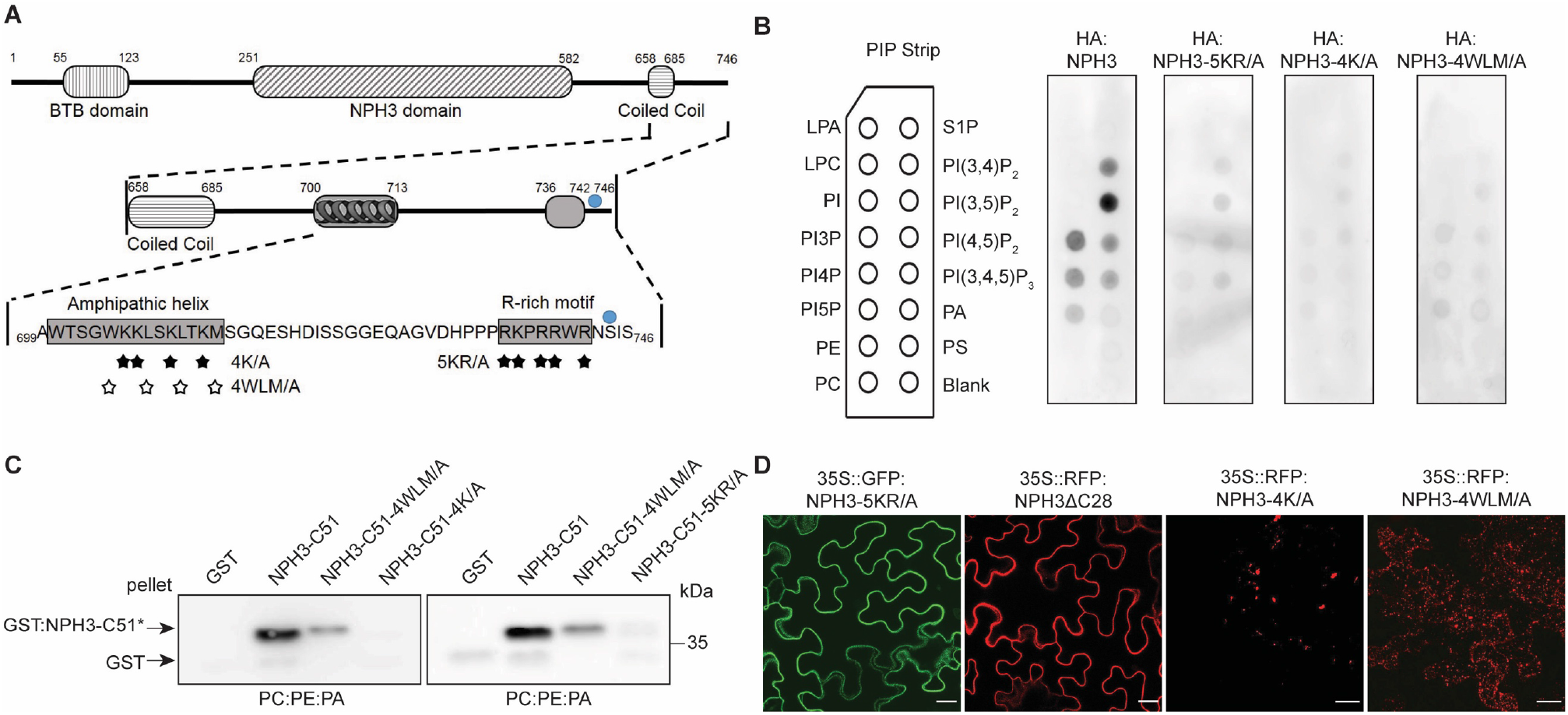
An amphipathic helix within the C-terminal domain is required for NPH3 phospholipid binding, membrane association and plasma membrane localization. **(A)** Domain structure and primary sequence of NPH3 showing the two putative BH domains (amphipathic helix and R-rich motif) within the C-terminal region. Stars depict residues of either the R-rich motif or the amphipathic helix substituted by alanine (A) in the NPH3 variants, blue circle depicts the 14-3-3 binding site. **(B)** Lipid overlay assay performed with purified GST:NPH3-C51 variants (C51*). **(C)** Liposome binding assay using large unilamellar liposomes containing the neutral PE and PC mixed with the polyacidic PA. Anti-GST immunoblot of GST:NPH3-C51 variants is shown. **(D)** Representative confocal microscopy images of leaf epidermal cells from transiently transformed *N. benthamiana* adapted to darkness (Z-stack projections of NPH3-4K/A and NPH3-4WLM/A are shown). Scale bars, 25 μm.

### NPH3 forms membrane-less condensates in the cytosol

As already mentioned, BL-induced PM dissociation and particle assembly of NPH3 in the cytosol seem to be separate and consecutive processes (Video S3). As yet, the identity of these particles has not been determined. NPH3ΔC51 is devoid of the amphipathic helix and localized to cytosolic particles in darkness (Fig. 3D). Subcellular fractionation clearly illustrated that the lack of the C-terminal region shifts NPH3 from a membrane-associated state to the soluble fraction (Fig. 3E). This reveals a non-membrane-attached state of NPH3 in discrete bodies as has been suggested for NPH3 aggregates generated upon BL irradiation [20]. Apparently, the mechanisms of NPH3 targeting towards and away from the PM are distinct from vesicle-mediated transport of transmembrane proteins. This is in line with the observation that NPH3 is insensitive to an inhibitor of endosomal trafficking [20]. Considering the lack of the 14-3-3 binding motif in NPH3ΔC51, 14-3-3 association seems dispensable for NPH3 body formation in the cytosol. To confirm this assumption, we examined NPH3 variants incapable of 14-3-3 binding, namely (i) NPH3-4K/A-S744A and (ii) NPH3-S744A, the latter upon co-expression with SAC1. Indeed, prevention of 14-3-3 association did not affect assembly of NPH3-4K/A-S744A particles in darkness (Fig. S4C). Moreover, similar to NPH3, NPH3-S744A localized to cytosolic particles in the dark upon co-expression of SAC1 but not SAC1DEAD (Fig. 3A). Hence, generation of NPH3 particles in the cytosol is feasible in the absence of 14-3-3s. It might thus be due to intrinsic properties of cytosolic NPH3 when exceeding a critical concentration. Taking constitutive PM association of NPH3-S744A in the absence of SAC1 into account, 14-3-3 association seems to be crucial for initial PM detachment while formation of discrete bodies occurs as an autonomous process in the cytosol.

The dynamic generation and morphology of NPH3 bodies is reminiscent of membrane-less biomolecular condensates which are micron-scale compartments in cells lacking surrounding membranes. An important organizing principle is liquid-liquid phase separation driven by multivalent macromolecular interactions – either mediated by modular interaction domains or disordered regions [40]. NPH3 is characterized by both intrinsically disordered regions and interaction domains such as the BTB and the CC domain (Fig. S1). We performed single-cell time-lapse imaging of RFP:NPH3 body formation to investigate whether NPH3 undergoes transition from a solute to a condensed state in *N. benthamiana*. Indeed, formation of particle-like structures in the cytosol is initiated after approx. 4 min and the fluorescence intensity per body gradually increased over time as a result of the growth in size (Fig. 2E, F). In contrast to the signal intensity, the number of bodies reached a maximum after approx. 10 to 15 min and afterwards started to decrease as a result of body fusion (Fig. 2E, G). Worth mentioning, these features are characteristic criteria of biomolecular condensates [40, 41].

### Phosphorylation of the 14-3-3 binding site in NPH3 is light-dependent and reversible

In dark-grown seedlings, NPH3 exists as a phosphorylated protein irrespective of phot1 activity. Light-induced dephosphorylation of NPH3 is almost a dogma in the literature. It has been recognized as a slight shift in electrophoretic mobility of NPH3 upon SDS-PAGE [18] and requires – in accordance with the light-induced formation of particle-like structures in the cytosol [20] – the photoreceptor phot1. In the following, (de)phosphorylation of NPH3, represented by a modification of its electrophoretic mobility, will be referred to as ‘general’ (de)phosphorylation of NPH3. Nonetheless, the data presented so far suggest that light-triggered and presumably S744 phosphorylation-dependent 14-3-3 association contributes to NPH3 function – an obvious antagonism to the ‘dogma of dephosphorylation’. Therefore, a phosphosite-specific peptide antibody (*α*-pS744) was established (antigen: _734_PPRKPRRWRN-S(P)-IS_746_) and an antibody against the unmodified peptide (*α*-NPH3) served as control. Examination of GFP:NPH3 in either *N. benthamiana* leaves or transgenic Arabidopsis lines revealed the typical enhanced electrophoretic mobility upon BL excitation, indicative of a ‘general’ dephosphorylation [18-20]. Intriguingly, the *α*-pS744 antibody recognized GFP:NPH3, but not GFP:NPH3-S744A, exclusively upon BL irradiation (Fig. 5; Fig. S5). Consequently, BL triggers two different posttranslational modifications of NPH3: (i) the phosphorylation of the 14-3-3 binding site (S744) and (ii) a ‘general’ dephosphorylation. Yet, neither of the modifications could be observed for GFP:NPH3-S744A (Fig. 5A, Fig. S5A). To uncover light-induced 14-3-3 association at the molecular level, an IP of GFP:NPH3 was conducted and combined with 14-3-3 Far Western analysis. Indeed, phosphorylation of S744 enabled binding of purified recombinant 14-3-3 proteins to NPH3 upon SDS PAGE (Fig. 5B, Fig. S5A). Prolonged irradiation or transfer of BL-irradiated seedlings to darkness is known to confer PM re-association of NPH3 [20], correlating with a reduced electrophoretic mobility indicative of a ‘general’ re-phosphorylation [18, 20]. Remarkably, we observed simultaneous dephosphorylation of S744 (Fig. 5B, Fig. S5B), effectively preventing binding of 14-3-3 to NPH3 (Fig. 5B). Taken together, the light/dark-dependent phosphorylation status of S744 determines 14-3-3 association with NPH3. Furthermore, the phosphorylation status of the 14-3-3 binding site and of NPH3 ‘in general’ is modulated by the light regime in an opposite manner, giving rise to a coinciding, but inverse pattern. Time course analyses, however, proved S744 phosphorylation of NPH3 to precede ‘general’ dephosphorylation upon BL treatment (Fig. S5B). In the past, ‘general’ dephosphorylation of NPH3 has been assumed to determine PM release of NPH3 coupled to particle assembly in the cytosol [4, 7, 12, 20]. Nonetheless, our data clearly indicated S744 phosphorylation-dependent 14-3-3 association to be the cause of PM dissociation, but not of condensate assembly in the cytosol. ‘General’ dephosphorylation might thus be coupled to PM dissociation/condensate formation. We examined the ‘general’ phosphorylation status of both NPH3 and NPH3-S744A when co-expressed with SAC1. Despite the fact that either NPH3 variant constitutively localized to cytosolic condensates (Fig. 3A), NPH3 was phosphorylated in darkness and shifted to the dephosphorylated status upon BL treatment while NPH3-S744A exhibited a permanent phosphorylated state (Fig. 5C). ‘General’ dephosphorylation of NPH3 is thus not coupled to PM dissociation. Furthermore, it is neither a prerequisite nor a consequence of condensate assembly, rather it seems to require prior light-triggered and S744 phosphorylation-dependent 14-3-3 association (Fig. 5A, C). Taken together, we suggest (Fig. 6E) that BL-induced and phosphorylation-dependent 14-3-3 association releases NPH3 from the PM into the cytosol and most likely enables ‘general’ dephosphorylation of NPH3. Formation of NPH3 condensates, however, is probably determined by the biological properties of PM-detached NPH3.

**Fig. 5:**
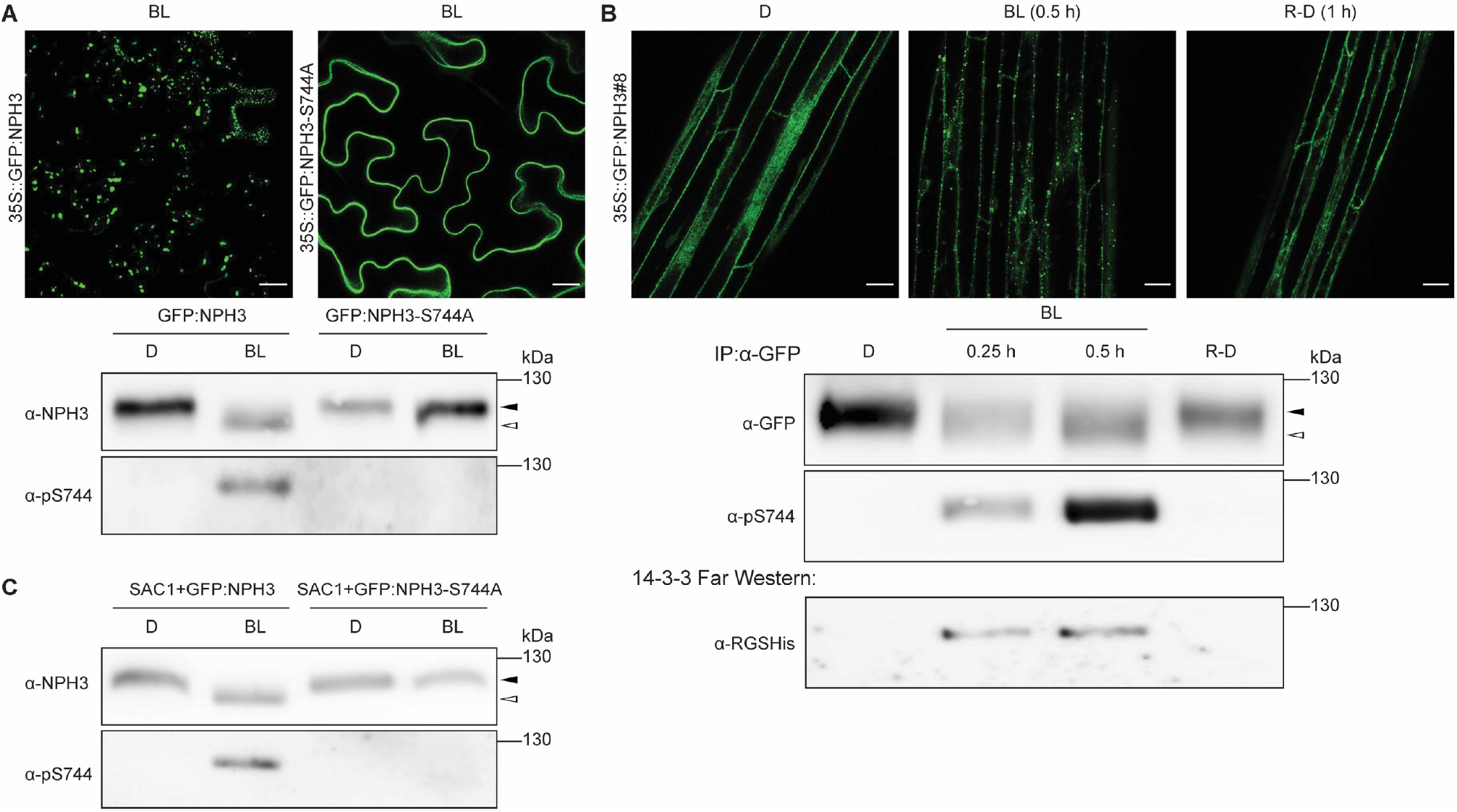
The phosphorylation status of the NPH3 14-3-3 binding site is dynamically modulated by the light regime. **(A)** Immunoblot analysis of total protein extracts from *N. benthamiana* leaves maintained in darkness (D) or treated with blue light (BL) (5 μmol m^-2^ sec^-1^) for 40 min (lower panel). The upper panel shows representative confocal microscopy images of leaf epidermal cells from transiently transformed *N. benthamiana* upon blue light treatment (40 min). Scale bars, 25 μm. **(B)** Immunoblot analysis and 14-3-3 Far-Western of anti-GFP immunoprecipitates from Arabidopsis *nph3-7* ectopically expressing GFP:NPH3. 3-days old etiolated seedlings were treated with cycloheximide (100 μM) for 1 h and either maintained in darkness (D), treated with blue light (BL) (1 μmol m^-2^ sec^-1^) for the indicated time, or re-transferred to darkness (1 h) after 30 min of irradiation (R-D). The immunoprecipitates were separated on 7.5% SDS-PAGE gels. The upper panel shows representative confocal microscopy images of hypocotyl cells from transgenic etiolated Arabidopsis seedlings under the specified conditions. Scale bars, 25 μm. **(C)** Immunoblot analysis of transiently transformed *N. benthamiana* leaves co-expressing SAC1:RFP with either GFP:NPH3 or GFP:NPH3-S744A and adapted to darkness (see Fig. 3A). Expression of transgenes was driven by the 35S promoter. Total protein extracts were separated on 7.5% SDS-PAGE gels. The closed and open arrowheads indicate the positions of ‘generally’ phosphorylated and dephosphorylated NPH3 proteins, respectively.

**Fig. 6:**
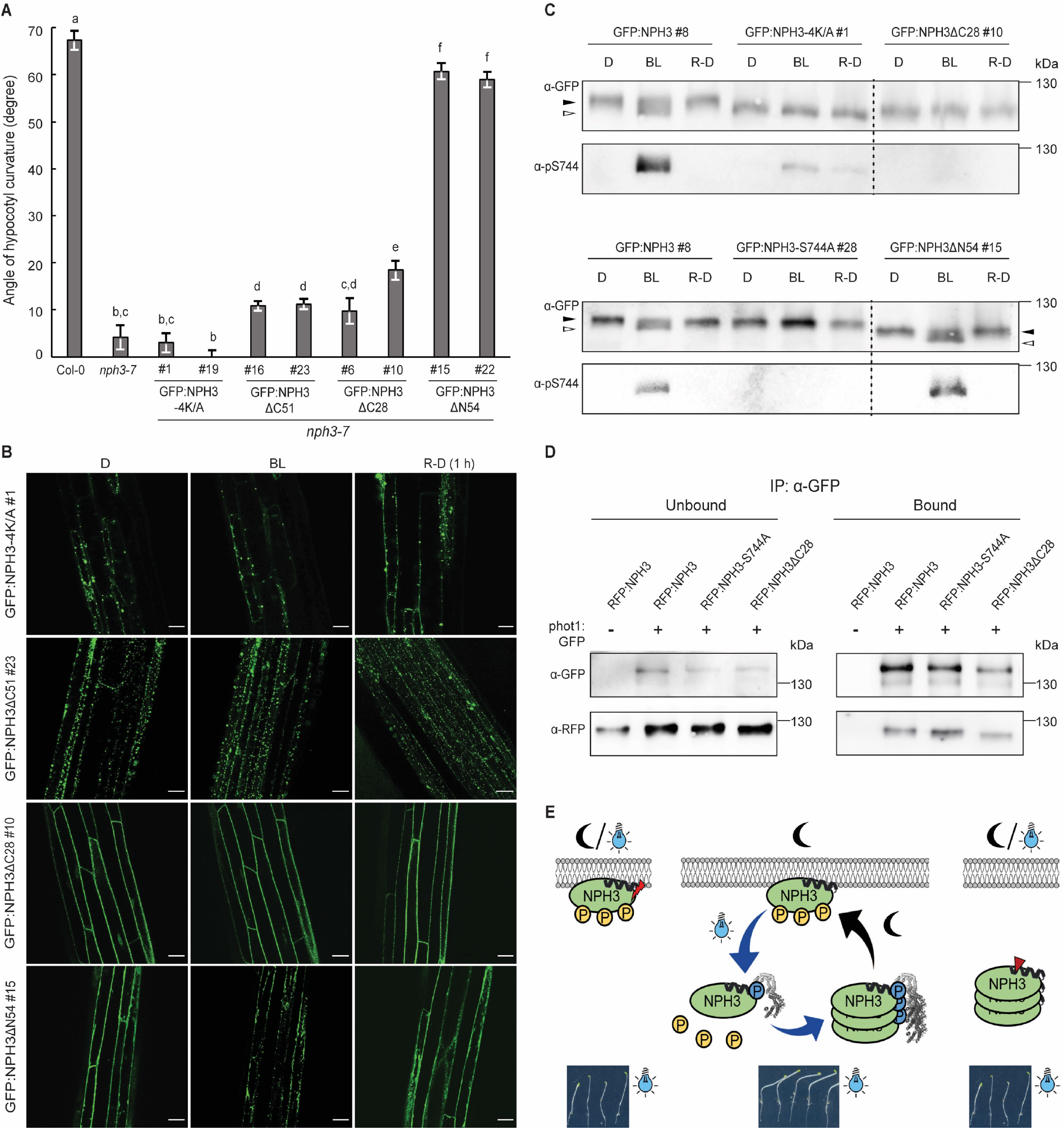
Functional relevance of the subcellular localization of NPH3. **(A)** Quantification of the hypocotyl phototropism response (mean ± SEM) in 3-days old etiolated seedlings exposed for 12 h to unilateral blue light (1 μmol m^-2^ sec^-1^) (n>30 seedlings per experiment, one representative experiment of two replicates is shown). Expression of wild-type and mutant variants of GFP:NPH3 in *nph3-7* was driven by the 35S promoter. Student’s t-test, different letters mark statistically significant differences (P<0.05), same letters mark statistically non-significant differences. **(B)** Representative confocal microscopy images of hypocotyl cells from transgenic Arabidopsis *nph3-7* seedlings ectopically expressing mutant variants of GFP:NPH3. 3-days old etiolated seedlings were either maintained in darkness (D), treated with blue light (BL) (approx. 11 min by means of the GFP-laser) or re-transferred to darkness (1 h) (R-D) after 30 min of irradiation (1 μmol m^-2^ sec^-1^). Scale bars, 25 μm. **(C)** Immunoblot analysis of etiolated Arabidopsis *nph3-7* seedlings ectopically expressing mutant variants of GFP:NPH3 and treated as described in **(B)**. Total protein extracts were separated on 7.5% SDS-PAGE gels. All samples shown in one panel are from the same blot, the dashed line was inserted to indicate an expected modification of the molecular weight of NPH3 due to deletions. The closed and open arrowheads indicate the positions of ‘generally’ phosphorylated and dephosphorylated NPH3 proteins, respectively. **(D)** *In vivo* interaction of RFP:NPH3 and phot1:GFP in transiently transformed *N. benthamiana* leaves adapted to darkness. Expression of transgenes was driven by the 35S promoter. Microsomal proteins were immunoprecipitated using GFP beads and separated on 11% SDS-PAGE gels, followed by immunoblotting with anti-GFP and anti-RFP antibodies, respectively. **(E)** Model depicting the light-regime triggered changes in the phosphorylation status, subcellular localization and phototropic responsiveness of NPH3. BL-induced and phosphorylation-dependent (S744, blue) binding of 14-3-3 proteins releases NPH3 from the PM into the cytosol followed by condensate formation. Residues that are phosphorylated in darkness (yellow) and become dephosphorylated upon light treatment give rise to a shift in electrophoretic mobility (‘general’ phosphorylation status). Re-transfer to darkness reverts all BL-triggered processes, finally resulting in PM re-association. Cycling of NPH3 between the PM and the cytosol seems to be essential for proper function. *Vice versa*, NPH3 variants either constitutively attached to (red flash) or constitutively detached (red arrowhead) from the PM are non-functional.

### Cycling of NPH3 might be key to function

The light-triggered and reversible shift in subcellular localization of NPH3 has led to the hypothesis that PM localization of NPH3 promotes its action in mediating phototropic signaling. In turn, NPH3 present in soluble condensates is considered to be inactive [7, 12, 20]. The functional relevance of the transient changes in subcellular NPH3 localization is, however, still not known. To assess the functionality of NPH3 variants constitutively localizing to condensates, GFP:NPH3-4K/A (Fig. 4D) as well as GFP:NPH3ΔC51 (Fig. 3D) were expressed in the *loss of function* Arabidopsis mutant *nph3-7*. Worth mentioning, the electrophoretic mobility of GFP:NPH3-4K/A corresponded to the dephosphorylated version of NPH3 and was not modified by light treatment (Fig. 6C). In line with the hypothesis mentioned above, NPH3 mutants constitutively present in condensates did not restore hypocotyl phototropism (Fig. 6A, B, Video S7, Video S8). Contrary to the hypothesis, however, GFP:NPH3-S744A - despite exhibiting constitutive PM localization (Fig. 2B-D) - is also largely incapable of mediating phototropic hypocotyl bending in *nph3-7* (Fig. 2A). To verify significantly impaired activity of permanently PM-attached NPH3, we examined NPH3ΔC28 in addition. Comparable to the results obtained in *N. benthamiana* (Fig. 4D), NPH3ΔC28 remained PM-associated upon activation of phot1 in stable transgenic Arabidopsis lines (Fig. 6B, Video S9) and its electrophoretic mobility was not modified by BL treatment (Fig. 6C). Noteworthy, both NPH3-S744A and NPH3ΔC28 still interacted with phot 1 (Fig. 6D), indicating that complex formation at the PM is not compromised. Nevertheless, permanent attachment of NPH3 to the PM turned out to be insufficient for triggering the phototropic response in *nph3-7* (Fig. 6A).

Taken together, neither NPH3 mutants permanently detached from the PM nor NPH3 versions permanently attached to the PM seem to be fully functional (Fig. 6A, E). So, what is the underlying mechanism of NPH3 function? We examined NPH3ΔN54 (Fig. 1A; Fig. 3D) in more detail. Similar to NPH3, NPH3ΔN54 associated to the PM in etiolated seedlings (Fig. 6B). Upon irradiation it (i) became phosphorylated at S744 (Fig. 6C), (ii) exhibited an increased electrophoretic mobility, indicative of a ‘general’ dephosphorylation (Fig. 6C) and (iii) detached from the PM followed by condensate formation in the cytosol (Fig. 6B, Video S10). Furthermore, all these processes were reverted when seedlings were re-transferred to darkness (Fig. 6B, C). Intriguingly, expression of NPH3ΔN54 completely restored phototropic hypocotyl bending in *nph3-7* (Fig. 6A) as did NPH3 (Fig. 2A). Thus, 14-3-3 mediated cycling of NPH3 between the PM and the cytosol might be of utmost importance for functionality (Fig. 6E).

## Discussion

Our data provide novel insight into the molecular mechanisms defining NPH3 function in BL-induced phototropic hypocotyl bending. We applied a combination of genetic, biochemical, physiological and live cell imaging approaches to uncover the impact of 14-3-3 proteins on NPH3, in particular its BL-triggered and phosphorylation-dependent PM release. Association of NPH3 with the PM is known since decades, but the underlying mechanism of membrane recruitment remained unknown. We demonstrated that NPH3 attaches to the PM in a phospholipid-dependent manner in darkness (Fig. 3A). The electrostatic interaction with polyacidic phospholipids (Fig. 3B, C) is mediated by four basic residues of an amphipathic helix, the hydrophobic face of which further contributes to PM association (Fig. 4D). We therefore suggest the amphipathic helix to be embedded in the PM inner-leaflet with its hydrophobic interface inserted in the hydrophobic core of the bilayer while the positively charged interface is arranged on the PM surface, interacting with the lipid polar heads. The amphipathic helix (amino acids 700-713) localizes downstream of the CC domain of NPH3 in its C-terminal region which also encompasses the 14-3-3 binding site (S744) (Fig. 1A, B). We discovered that BL induces two different, but coinciding posttranslational modifications in NPH3 (Fig. 5): (i) the phosphorylation of S744 which in turn enables association of 14-3-3 proteins with NPH3 and (ii) the well-described dephosphorylation, represented by an enhanced electrophoretic mobility of NPH3 (‘general’ dephosphorylation) [18-20]. The - as yet unrecognized - BL-induced phosphorylation event linked to 14-3-3 association is of utmost importance since it is essential for (i) the BL-triggered internalization of NPH3 from the PM (Fig. 2B) and (ii) the function of NPH3 in phototropic hypocotyl bending (Fig. 2A).

However, expression of NPH3-S744A which is incapable of 14-3-3 interaction, partially restored the severe impairment of hypocotyl phototropism in *nph3-7* (Fig. 2A). This might be due to functional redundancy among certain members of the NRL protein family. Indeed, RPT2, expression of which is induced and stabilized by BL treatment [42], is required for hypocotyl phototropism at light intensities utilized in our assays [20]. RPT2 might therefore – at least in part – substitute for NPH3. The same applies to DEFECTIVELY ORGANIZED TRIBUTARIES 3 (DOT3), the, as yet, functionally uncharacterized closest homolog of NPH3 [12]. Worth mentioning, RPT2, DOT3 and also the NRL protein MACCHI-BOU 4 (MAB4)/ ENHANCER OF PINOID (ENP) are capable of interacting with 14-3-3 proteins in yeast via their antepenultimate serine residues, respectively (our unpublished data). Alternatively, residual activity of NPH3-S744A in phototropic hypocotyl bending might be caused by its permanent association with the PM. Light treatment could induce a reorganization of NPH3-S744A within/along the PM which might allow for phototropic responsiveness to a certain level. Addressing these alternatives represents a formidable challenge for future research. NPH3 has been described to re-localize directly from the PM into discrete bodies in the cytosol upon light treatment [7, 20]. It became, however, evident that it initially detaches from the PM into the cytosol (Video S3). Here, NPH3 undergoes a dynamic transition from a dilute to a condensed state, resulting in the formation of membrane-less biomolecular compartments (Fig. 2E; Fig. 3E). Biomolecular condensates are emerging as an important concept in signaling, also in plants [43]. Their formation can be driven by multivalent interactions with other macromolecules, by intrinsically disordered regions within a single molecule or both [40, 44]. Interestingly, 14-3-3 proteins are dispensable for condensate assembly in the cytosol, as demonstrated by diverse NPH3 mutant variants (Fig. 3A, D; Fig. S4C). Further studies will reveal whether condensate formation of the PM-detached NPH3 is essential for its action.

As described above, the light-triggered modifications of the phosphorylation pattern of NPH3 are highly complex. Our observations disproved the view that BL-triggered ‘general’ dephosphorylation events determine PM dissociation of NPH3 [7, 12, 20]. First of all, dephosphorylation of NPH3 – i.e. a decrease in negative charge - is entirely inappropriate to interfere with membrane association relying on electrostatic interactions with polyacidic phospholipids (Fig. 3B, Fig. 4D). Furthermore, investigation of the seven NPH3 phosphorylation sites that were recently identified in etiolated Arabidopsis seedlings revealed that the phosphorylation status of these NPH3 residues was neither required for PM association in darkness nor BL-induced release of NPH3 into the cytosol [21]. By contrast, single site mutation of the 14-3-3 binding site in NPH3 (S744A) abolished PM dissociation upon BL treatment (Fig. 2B-D), indicating light-induced and phosphorylation-dependent 14-3-3 association to mediate PM release of NPH3. Given that the amphipathic helix localizes approximately 30 – 45 residues upstream of the 14-3-3 binding site (Fig. 4A), 14-3-3 binding to NPH3 is expected to induce a substantial conformational change that liberates the amphipathic helix from the PM. The molecular mechanism of NPH3 internalization is hence different from the - likewise PM-associated - photoreceptor phot1, trafficking of which occurs via vesicles through the endosomal recycling pathway [45]. Now, what about the BL-triggered ‘general’ dephosphorylation of NPH3? Based on our findings, this posttranslational modification temporally succeeded light-induced S744 phosphorylation (Fig. S5B). Furthermore, ‘general’ dephosphorylation was coupled to BL-triggered S744 phosphorylation, irrespective of the subcellular localization of NPH3 (Fig. 5A, C). We therefore assume phosphorylation-dependent 14-3-3 binding to be required for BL-induced ‘general’ dephosphorylation of NPH3 as well - a hypothesis that will be examined by future research.

Re-transfer of BL-irradiated seedlings to darkness triggers (i) dephosphorylation of S744 linked to 14-3-3 dissociation. 14-3-3 release is expected to result in a re-exposure of the amphipathic helix, which subsequently mediates (ii) re-association with the PM and presumably (iii) re-phosphorylation of NPH3, represented by a reduced electrophoretic mobility (‘general’ re-phosphorylation) (Fig. 5B, Fig. S5B). Intriguingly, neither NPH3 variants that constitutively localize to the PM nor mutant versions constitutively detached from the PM are capable of restoring the severe defect in hypocotyl phototropism in *nph3-7*. Complementation of the *nph3-7* phenotype exclusively could be observed upon expression of NPH3 variants that exhibit a light regime-driven dynamic change in subcellular localization (Fig. 6A, B, C). In summary, we propose a model where S744 phosphorylation-dependent and 14-3-3 driven cycling of NPH3 between the PM and the cytosol critically determine NPH3 function in mediating phototropic signaling in Arabidopsis (Fig. 6E).

In the past, it has been hypothesized that the light-induced internalization of phot1 – first described in 2002 [46] - may be coupled to light-triggered re-localization of auxin transporters. Functionality of phot1, however, was unaffected when internalization of the photoreceptor was effectively prevented by PM tethering via lipid anchoring [47]. Altogether, the change in subcellular localization does not seem to be essential for signaling of phot1, but of its downstream signaling component NPH3 (Fig. 6E). Plant 14-3-3 proteins have been shown to contribute to the subcellular polar localization of PIN auxin efflux carrier and consequently auxin transport-dependent growth [23]. NRL proteins in turn act as signal transducers in processes involving auxin (re)distribution in response to developmental or environmental signals [23], hence providing a likely link between 14-3-3 and PIN polarity. One subfamily of the NRL protein family consists of MAB4/ ENP-like (MEL) polypeptides, playing a critical role in auxin-regulated organogenesis in Arabidopsis [48-50]. MEL proteins exhibited a polar localization at the cell periphery which was almost identical to that of PIN proteins [51, 52]. Remarkably, MEL proteins were recently shown to maintain PIN polarity by limiting lateral diffusion [53]. Thus, one attractive hypothesis is that certain NRL proteins contribute either to the maintenance or to a dynamic change of the subcellular polarity of PIN auxin carriers, thereby regulating auxin (re)distribution. With respect to hypocotyl phototropism, light-induced and 14-3-3-mediated detachment of NPH3 from the PM might account for BL-driven changes in PIN polarity. Future studies will address the impact of NPH3 on the subcellular polarity of PIN proteins.

## Material and Methods

### Plant materials, transformation and growth conditions

*Arabidopsis thaliana* (ecotype Columbia-0 (Col-0)) expressing 14-3-3 epsilon:GFP under control of the native promoter has been described recently [23]. Seeds of *A. thaliana nph3-7* (SALK_110039, Col-0 background) were obtained from the Nottingham Arabidopsis Stock Centre. T-DNA insertion was confirmed by genomic PCR analysis and homozygous lines were identified. Stable transformation of *nph3-7* followed standard procedures. Seeds were surface sterilized and planted on solid half-strength Murashige and Skoog (MS) medium (pH 5.8). Following stratification in the dark for 48-72 h at 4°C, seeds were exposed to fluorescent white light for 4 h. Subsequently, seedlings were grown at 20°C in darkness for 68 h. Light treatment of etiolated seedlings was done as specified in the Figure legends. Independent experiments were carried out at least in triplicates with the same significant results. Representative images are presented. Statistics were evaluated with Excel (Microsoft).

Transient transformation of 3-4 weeks old *Nicotiana benthamiana* plants was performed exactly as described [54]. Freshly transformed tobacco plants were kept under constant light for 24 h, subsequently transferred to darkness for 17 h (dark adaptation) and finally irradiated or kept in darkness as specified in the Figure legends.

### Cloning procedures

A 2.1 kb *NPH3* promoter fragment was PCR-amplified from Col-0 genomic DNA and the cDNA of NPH3 was amplified from Col-0 cDNA. The respective primers were characterized by BsaI restriction sites allowing for the usage of the Golden Gate based modular assembly of synthetic genes for transgene expression in plants [55]. Following A-tailing, the individual PCR products were directly ligated into the pGEM-T Easy (Promega) vector yielding level I vectors LI A-B p*NPH3* and LI C-D NPH3, respectively. Golden Gate level II assembly was performed by BsaI cut ligation and by using the modules LI A-B p*NPH3*, LI B-C GFP or LI B-C mCherry, LI C-D NPH3, LI dy D-E, LI E-F nos-T and LI F-G Hygro exactly as described [55].

For CoIP of fluorophore-tagged NPH3 and 14-3-3 transiently expressed in *N. benthamiana*, the corresponding cDNA was cloned into the 2in1 GATEWAY™ compatible vector pFRETcg-2in1-NC [56] via GATEWAY™ technology.

Cloning of N-terminally fluorophore-tagged NPH3 variants (GFP and/or RFP) into the destination vectors pB7WGR2 and/or pH7WGF2 [57] for stable or transient overexpression followed standard GATEWAY™ procedures. Transgenic plants were selected based on the hygromycin resistance conferred by pH7WGF2 and homozygous lines were established. The *35S*-driven *PHOT1:GFP* [45] and the 35S::MAP:mCherry:SAC1/SAC1_DEAD_ transformation vectors [33] as well as the utilized Golden Gate level I vectors [55] have been described before, respectively.

Site-directed mutagenesis was performed by PCR. PCR products and products of mutagenesis were verified by sequencing.

A complete list of oligonucleotides used for PCR is provided below.

### Expression and purification of proteins

For expression of the Arabidopsis 14-3-3 isoform omega as RGS(His)_6_-tagged protein in *Escherichia coli* M15, the corresponding cDNA was amplified by PCR and cloned into the expression vector pQE-30 (Qiagen). Purification was done by using Ni^2+^-NTA agarose (Qiagen) according to the manufacturer’s protocol.

For expression of the Arabidopsis NPH3 C-terminal 51 residues fused to GST in *E. coli* BL21(DE3), the corresponding cDNA fragment was amplified by PCR and cloned into the GST expression vector pGEX-4T-1. GST fusion proteins were purified from transformed bacteria using GSH-Sepharose according to the manufacturer’s protocol (Cytiva). Free GST protein was expressed and purified as a negative control.

### Cell-free protein expression

Reactions were performed using the TNT® T7 Quick Coupled Transcription/Translation System (Promega) with 1 μg of vector (NPH3 or variants in pGADT7) for a 50 μl reaction. Protein expression was carried out at 30°C for 90 min. Immunodetection was performed by using an anti-HA antibody (HA-tag encoded by pGADT7).

### Preparation of microsomal membranes

Microsomal membrane fractions were prepared from transiently transformed *N. benthamiana* leaves. Tissue was homogenized with 3 mL homogenization buffer per g fresh weight (50 mM Hepes (pH 7.8), 500 mM sucrose, 1 % (w/v) PVP-40, 3 mM DTT, 3 mM EDTA, supplemented with Complete Protease Inhibitor Mixture (Roche) and Phosphatase Inhibitor Mix 1 (Serva)). The homogenate was centrifuged at 10,000 g for 20 min at 4 °C. The supernatant was filtered through MiraCloth and subsequently centrifuged at 100,000 g for 45 min at 4 °C. The microsomal pellet was resuspended in 5 mM Tris/MES (pH 6.5), 330 mM sucrose, 2 mM DTT, supplemented with Complete Protease Inhibitor Mixture (Roche) and Phosphatase Inhibitor Mix 1 (Serva).

### Phospholipid binding assays

For lipid binding assays, either NPH3 variants expressed in a cell free system or purified recombinant GST fusion proteins were applied. Lipid overlay assays using PIP-strips were performed following the manufacturer’s instructions (Echelon). In brief, membranes were blocked overnight at 4°C in a blocking buffer with 4% fatty acid-free BSA in PBS-T (0.1% Tween). Purified proteins (0.1 μg/ml blocking buffer) or 10-50 μl of the cell free expression reaction (volume adjusted according to prior immunodetection of individual reactions) were incubated with PIP-strip membranes for 1 h at room temperature and washed three times for 10 min with PBS-T. Subsequently, detection of bound proteins was done by immunodetection of either GST (GST fusion proteins) or the HA-tag (cell free expression). Liposome binding assays were conducted essentially as described by [58] with slight modifications. All lipids were obtained from Avanti Polar Lipids. Liposomes were prepared from 400 nmol of total lipids at the following molar ratios: PC:PE, 1:1; PC: PE:PI4P, 2:2:1; PC:PE:PA, 2:2:1. The binding buffer (150 mM KCl, 25 mM Tris–HCl pH 7.5, 1 mM DTT, 0.5 mM EDTA) was supplemented with Complete Protease Inhibitor Mixture (Roche). Purified GST-NPH3-C51 variants in binding buffer were centrifuged at 50,000 g to get rid of any possible precipitates. Following incubation of liposomes and proteins, the liposome pellet was washed twice with binding buffer. Liposome-bound GST-NPH3-C51 variants were detected by immunoblotting with anti-GST antibodies.

### Y2H, SDS-PAGE and Western Blotting

For yeast two-hybrid analyses, the individual constructs were cloned into the vectors pGADT7 and pGBKT7 and co-transformed into the yeast strain PJ69-4A. Activity of the *ADE2* reporter was analyzed by growth of co-transformed yeast on SD medium lacking adenine.

SDS-PAGE, Western blotting and immunodetection followed standard procedures. Total proteins were extracted from 3-day-old etiolated Arabidopsis seedlings (50 seedlings) or transiently transformed *N. benthamiana* leaves (2 leaf disks) by directly grinding in 100 μl 2 x SDS sample buffer under red safe light illumination. Chemiluminescence detection was performed with an Amersham Image Quant800 (Cytiva) system.

The rabbit anti-NPH3-S744P antibody was generated with the phosphorylated synthetic peptide NH_2_-PPRKPRRWRN-S(PO_3_H_2_)-IS-COOH followed by affinity-purifications against the non-phosphorylated and phosphorylated peptide at Eurogentec (Liege, Belgium).

### CoIP and mass spectrometry analysis

Arabidopsis seedlings expressing 14-3-3 epsilon-GFP (endogenous promoter) and, as control, GFP (UBQ10 promoter) were grown in the dark on half-strength MS plates for 3 days. Subsequently, the etiolated seedlings were either kept in darkness or treated with overhead BL (1 μmol m^-2^ sec^-1^) for 30 min. Three grams of plant tissue were used under red safe light illumination for immunoprecipitation as described [59]. The final precipitate in Laemmli buffer was analyzed by mass spectrometry (MS) at the University of Tübingen Proteome Center. Following a tryptic in gel digestion, LC-MS/MS analysis was performed on a Proxeon Easy-nLC coupled to an QExactiveHF mass spectrometer (method: 60 min, Top7, HCD). Processing of the data was conducted using MaxQuant software (vs 1.5.2.8). The spectra were searched against an *Arabidopsis thaliana* UniProt database. Raw data processing was done with 1% false discovery rate setting.

Two individual biological replicates were performed and the following candidates were omitted from the list of epsilon-GFP interaction partners: (i) proteins that interacted with GFP (control), (ii) proteins that were identified in only one of the two experiments. Protein signal intensities of well-known 14-3-3 client proteins (Fig. 1) were converted to normalized abundance of the bait protein. Fold changes in relative abundance of BL treatment versus darkness (BL *vs*. D) were calculated (Table S1).

Arabidopsis *nph3-7* ectopically expressing GFP:NPH3 and *N. benthamiana* leaves transiently overexpressing fluorophore-tagged proteins were immunoprecipitated under red safe light illumination according to [60]. Growth and light irradiation of the plants is specified elsewhere.

*In vivo* interaction of phot1:GFP and N-terminally RFP-tagged NPH3 variants was tested by using solubilized microsomal proteins obtained from dark adapted *N. benthamiana* plants ectopically co-expressing the proteins of interest. Solubilization was achieved by adding 0,5% Triton X-100 to resuspended microsomal proteins followed by centrifugation at 50,000 g for 30 min at 4 °C. The supernatant was added to GFP-Trap Beads (ChromoTek) and incubated at 4°C for 1 h. Precipitated beads were washed six times with 50mM HEPES pH 7.8, 150mM NaCl, 0,2% Triton X-100. Finally, proteins were eluted by SDS sample buffer and separated by SDS-PAGE.

### 14-3-3 Far-Western

Anti-GFP immunoprecipitates obtained from Arabidopsis *nph3-7* stably overexpressing GFP:NPH3 were separated by SDS-PAGE and transferred to nitrocellulose. Nonspecific sites were blocked by incubation with 4% (w/v) milk powder in TBS at room temperature for at least 1 h. Subsequently, the membrane was incubated overnight at 4 °C (followed by 1 h at room temperature) with purified recombinant RGS(His)_6_-tagged 14-3-3 isoform omega of Arabidopsis diluted to 20 μg ml^-1^ in 50 mM MOPS/NaOH, pH 6.5, 20% (w/v) glycerol, 5mM MgCl_2_, and 2mM DTT. After washing with TBS, immunodetection of RGS(His)_6_-tagged 14-3-3 was performed by applying the anti-RGS(His)_6_ antibody (Qiagen) in combination with a secondary anti-mouse HRP antibody.

### Hypocotyl Phototropism analysis

*A. thaliana* seedlings were grown in the dark on vertically oriented half-strength MS plates for 48 h. Etiolated seedlings were then transferred to a LED chamber and illuminated with unilateral BL (1 μmol m^-2^ sec^-1^) for 24 h. Plates were scanned and the inner hypocotyl angle was measured for each seedling using Fiji. The curvature angle was calculated as the difference between 180° and the measured value.

### Confocal microscopy

Live-cell imaging was performed using the Leica TCS SP8 (upright) confocal laser scanning microscope. For excitation and emission of fluorophores, the following laser settings were used: GFP, excitation 488 nm, emission 505-530 nm; RFP, excitation 558 nm, emission 600-630 nm. All CLSM images in a single experiment were captured with the same settings using the Leica Confocal Software. All the experiments were repeated at least three times. Images were processed using LAS X light.

Single-cell time-lapse imaging was carried out on live leaf tissue samples from *N. benthamiana* transiently expressing RFP:NPH3. PM-detachment was induced by means of the GFP-laser (488 nM) and image acquisition (RFP-laser) was done for the duration of 32 min by scanning 30 consecutive planes along the Z axis covering the entire thickness of an epidermal cell. Z-projection was done for each 3,5 min interval. For all image quantifications, randomly sampled unsaturated confocal images (512 x 512 pixels, 225 x 225 μm) were used with an image analysis protocol implemented in the ImageJ software [61] as previously described [62]. A random image was selected from the dataset and parameters such as local threshold, background noise, object size and shape were determined. The obtained parameters were used for image analysis of the whole dataset following exactly the published step by step protocol [62].

### List of primers used in this study

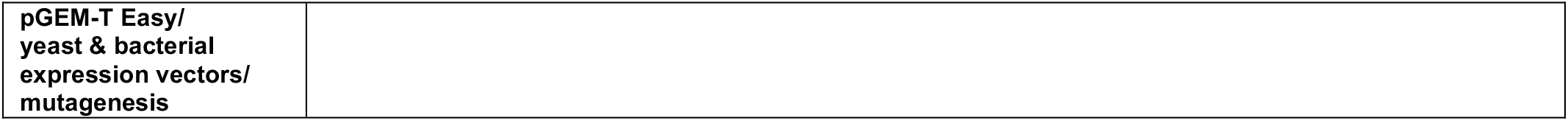
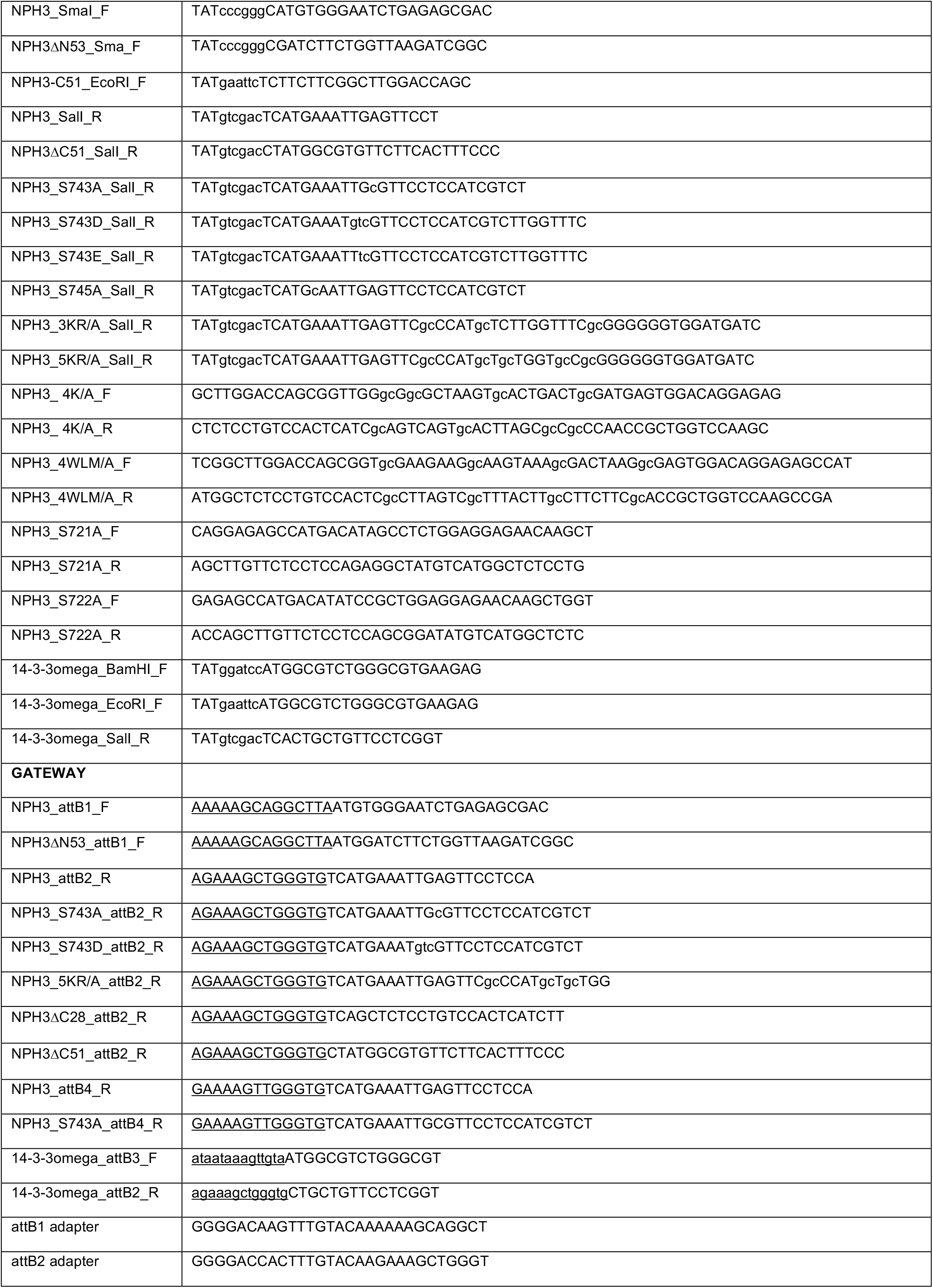
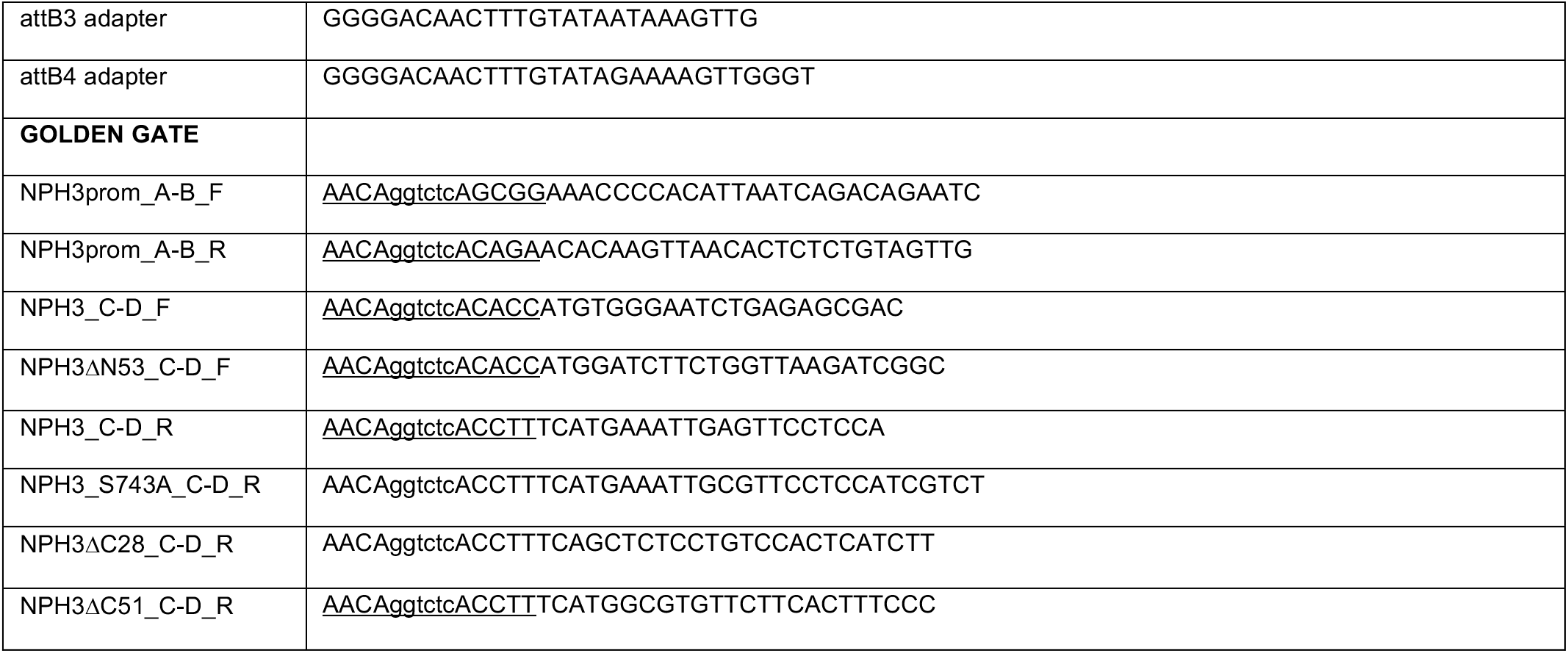

## Acknowledgements

We are grateful to Yvon Jaillais and John M. Christie for providing the constructs 35S::MAP:SAC1/SAC1_DEAD_ and 35S::PHOT1:GFP, respectively. We furthermore thank Sandra Richter for SP8 support and John M. Christie for stimulating discussions. MS analysis was done at the Proteome Centre, University of Tübingen, and we thank Irina Droste-Borel for help in data assessment. Research in our laboratory was supported by the German Research Foundation (DFG) with a grant to C.O. (CRC 1101-B09).

## Figures, tables, videos

**Fig. S1:**
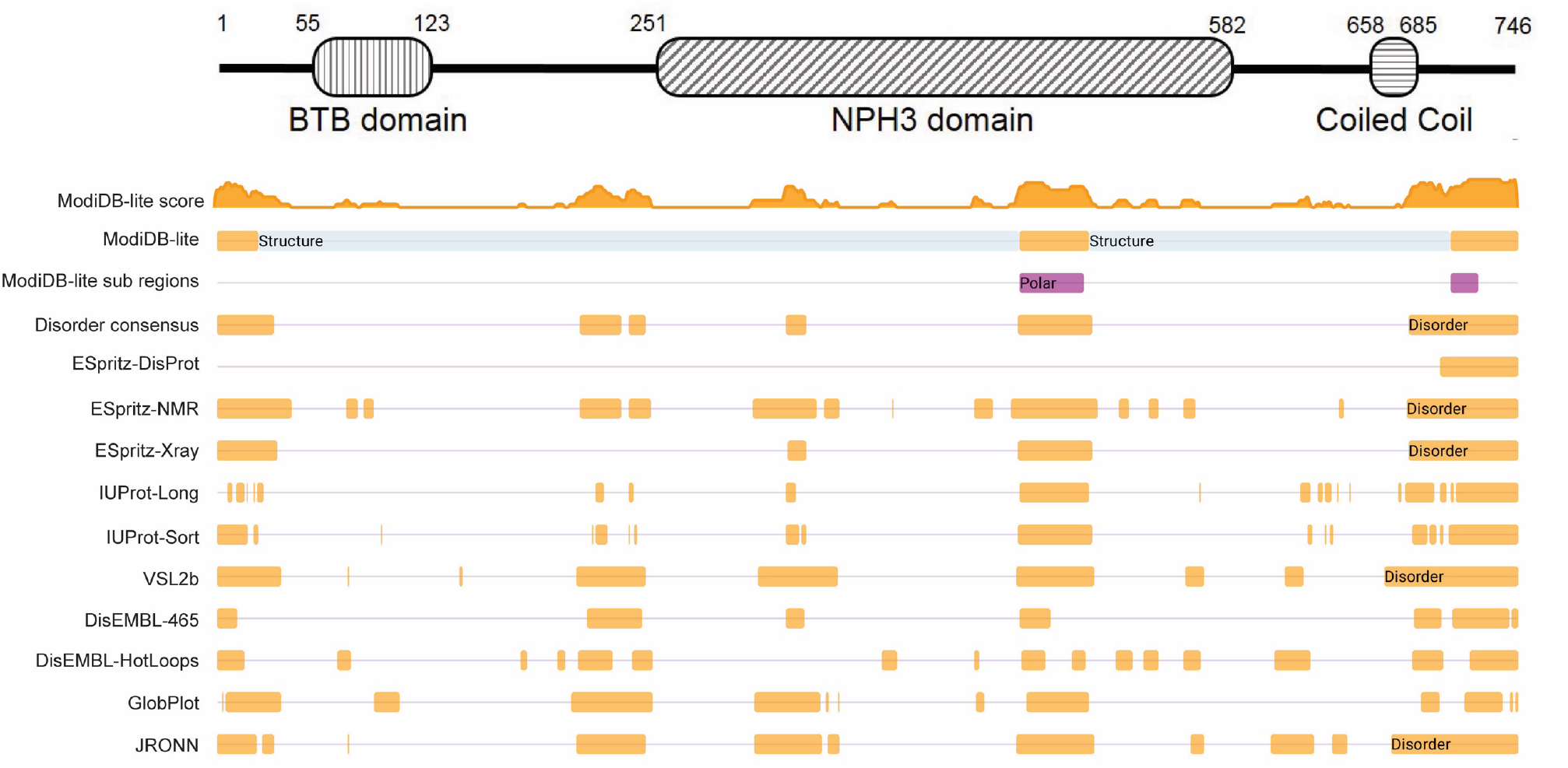
Domain structure of NPH3 and MobiDB plot (https://mobidb.org/) of intrinsically disordered regions in NPH3. BTB domain, <u>b</u>road-complex, tramtrack, <u>b</u>ric a brac domain.

**Table S1:**
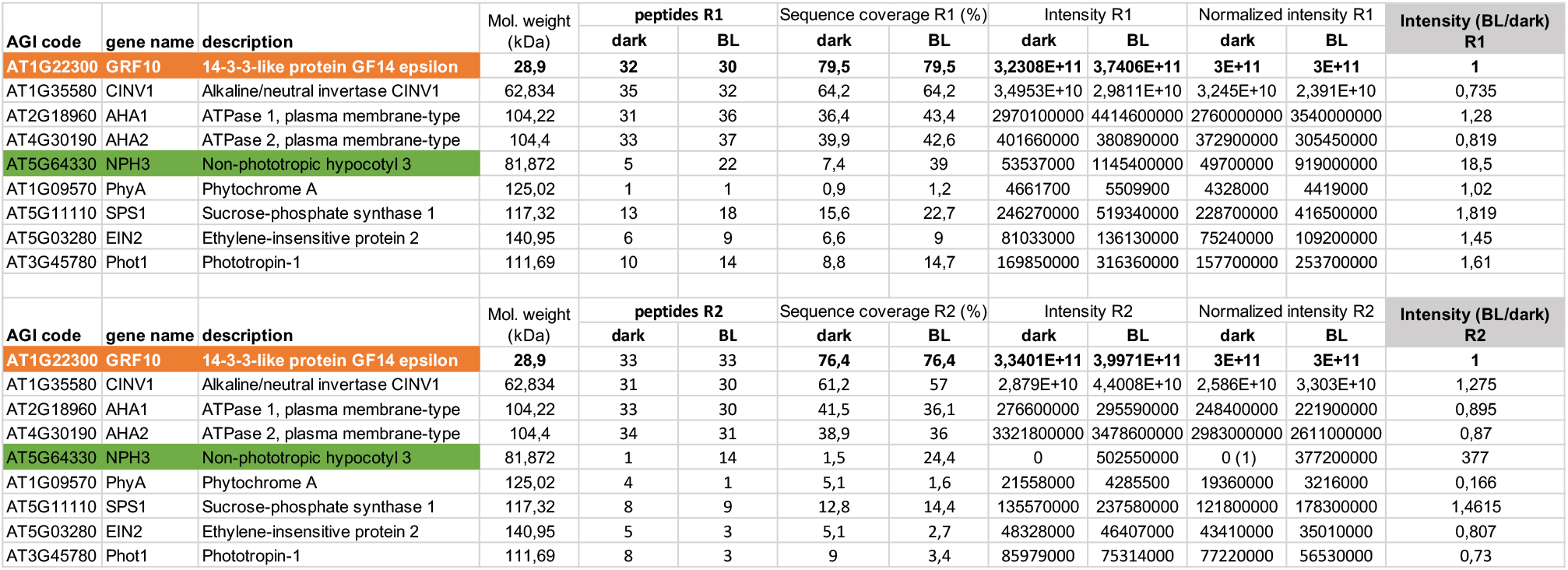
Analysis of 14-3-3 epsilon-GFP immunoprecipitates via mass spectrometry (MS) based on two biological replicates. This table lists only known 14-3-3 clients in addition to NPH3.

**Fig. S2:**
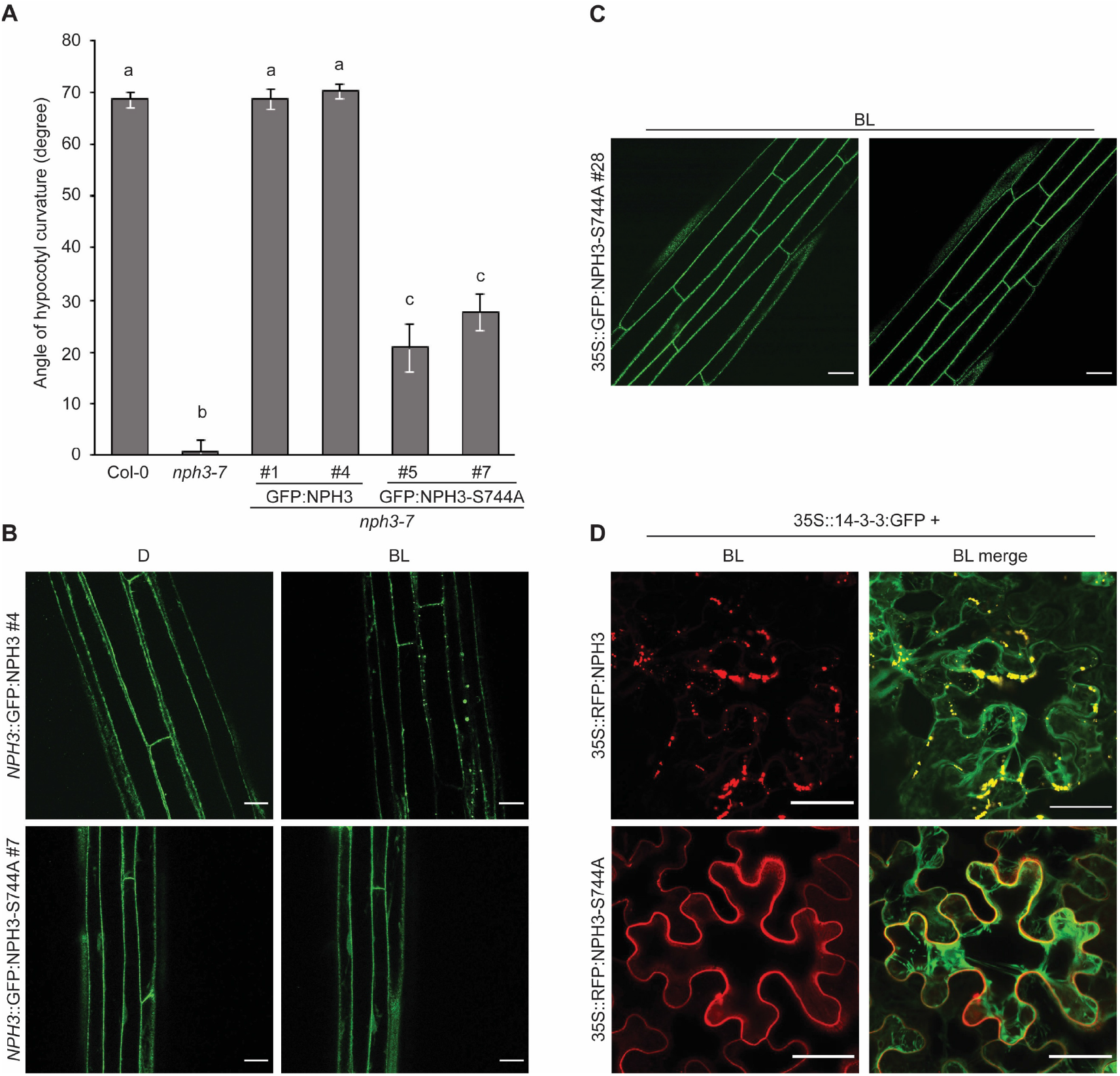
**(A)** Quantification of the hypocotyl phototropism response (mean ± SEM) in 3-days old etiolated seedlings exposed for 12h to unilateral blue light (1 μmol m^-2^ sec^-1^) (n>30 seedlings per experiment, one representative experiment of two replicates is shown). Expression of transgenes in *nph3-7* was driven by the *NPH3* promoter. Student’s t-test, different letters mark statistically significant differences (P<0.05), same letters mark statistically non-significant differences. **(B, C, D)** Representative confocal microscopy images of hypocotyl cells from transgenic etiolated Arabidopsis *nph3-7* seedlings **(B, C)** or of leaf epidermal cells from transiently transformed *N. benthamiana* (here, Z-stack projections are shown) **(D)**. The plants were either kept in darkness (D) or treated with blue light (BL) (*nph3-7*: 1 μmol m^-2^ sec^-1^ and *N. benthamiana*: 10 μmol m^-2^ sec^-1^) for 40 min. Expression of transgenes was driven by the *NPH3* promoter **(B)** or the 35S promoter **(C, D)**. Scale bars, 25 μm.

**Video S1**:

Dynamic BL-induced changes in the subcellular localization of 35S::GFP:NPH3 in hypocotyl cells of stably transformed Arabidopsis *nph3-7*.

**Video S2**:

Subcellular localization of 35S::GFP:NPH3-S744A in hypocotyl cells of stably transformed Arabidopsis *nph3-7* upon BL-irradiation.

**Video S3**:

Dynamic BL-induced changes in the subcellular localization of 35S::RFP:NPH3 transiently expressed in *N. benthamiana* leaves.

**Video S4**:

Subcellular localization of 35S::RFP:NPH3-S744A in transiently transformed *N. benthamiana* leaves upon BL-irradiation.

**Fig. S3:**
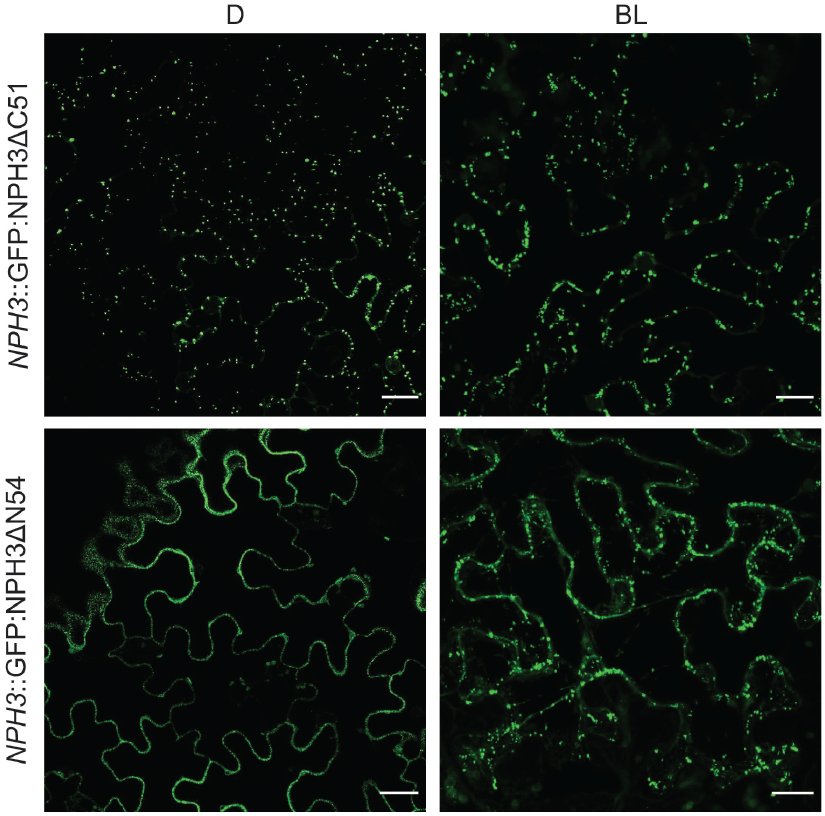
Representative confocal microscopy images of leaf epidermal cells from transiently transformed *N. benthamiana* (Z-stack projections of BL-treated NPH3 variants are shown). The plants were either kept in darkness (D) or treated with blue light (BL) (approx.11 min by means of the GFP-laser). Scale bars, 25 μm.

**Video S5**:

Subcellular localization of 35S::RFP:NPH3ΔC51 in transiently transformed *N. benthamiana* leaves upon BL-irradiation.

**Video S6**:

Dynamic BL-induced changes in the subcellular localization of 35S::RFP:NPH3ΔN54 transiently expressed in *N. benthamiana* leaves.

**Fig. S4:**
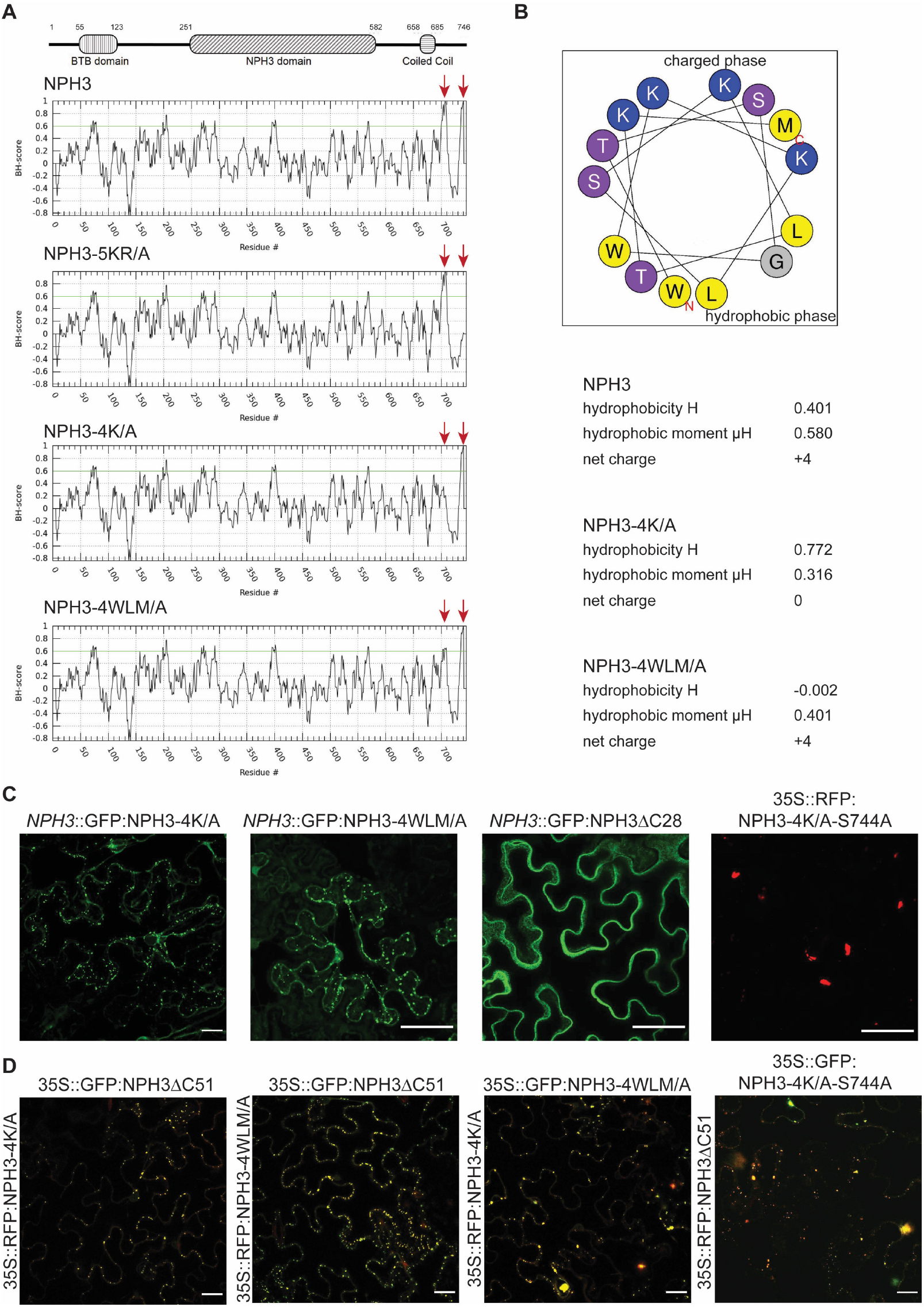
**(A)** BH score profiles (window size 11) of NPH3 and mutant variants. Putative BH-domains are indicated by red arrows. **(B)** Helical wheel projection showing amphipathy of the predicted helix (residues 700-713) within the C-terminal domain of NPH3. Overall helix hydrophobicity (H) and the hydrophobic moment (μH) are given for NPH3 and mutant variants. **(C)** Representative confocal microscopy images (Z-stack projections) of leaf epidermal cells from transiently transformed *N. benthamiana* adapted to darkness. Scale bars, 25 μm. **(D)** Representative merge fluorescence images of leaf epidermal cells from *N. benthamiana* transiently co-expressing NPH3 variants and adapted to darkness. Scale bars, 25 μm.

**Fig. S5:**
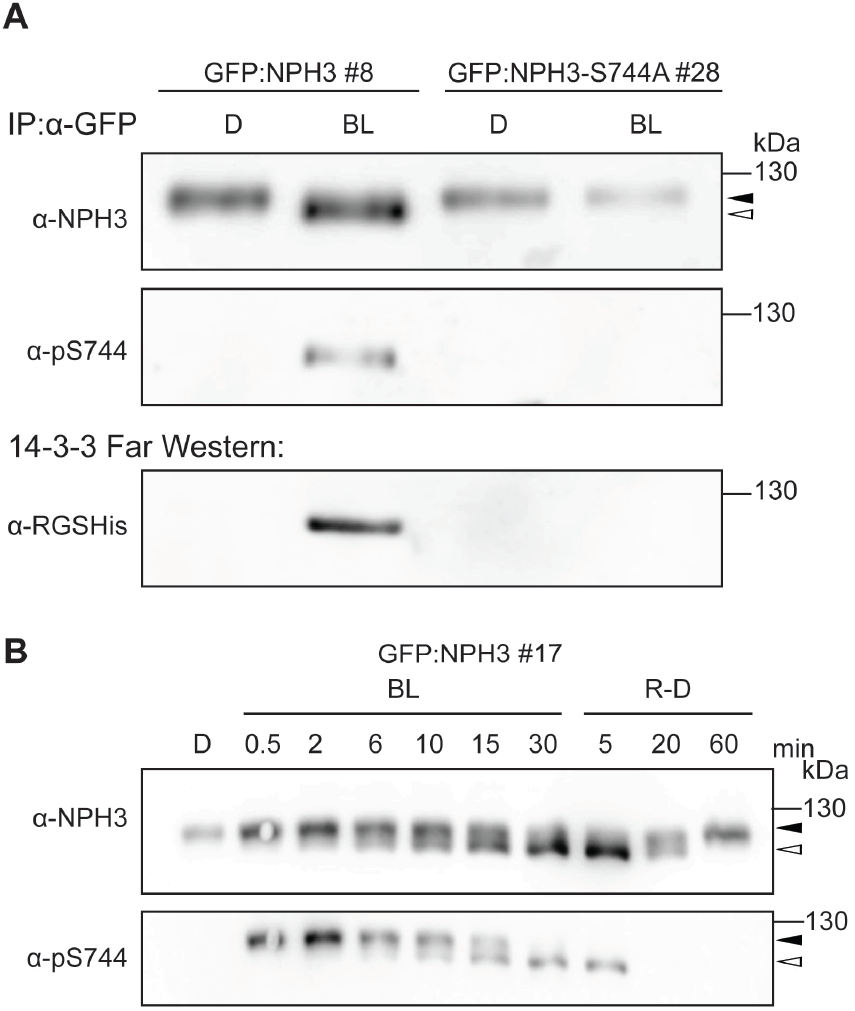
**(A)** Immunoblot analysis and 14-3-3 Far-Western of anti-GFP immunoprecipitates from Arabidopsis *nph3-7* ectopically expressing GFP:NPH3 or GFP:NPH3-S744A. 3-days old etiolated seedlings were either maintained in darkness (D) or treated with blue light (BL) (1 μmol m^-2^ sec^-1^) for 40 min. The immunoprecipitates were separated on 7.5% SDS-PAGE gels. **(B)** Immunoblot analysis of Arabidopsis *nph3-7* ectopically expressing GFP:NPH3. 3-days old etiolated seedlings were either maintained in darkness (D), treated with blue light (BL) (1 μmol m^-2^ sec^-1^) for the indicated time or re-transferred to darkness for the indicated time after 30 min of irradiation (R-D). Total protein extracts were separated on 7.5% SDS-PAGE gels. The closed and open arrowheads indicate the positions of ‘generally’ phosphorylated and dephosphorylated NPH3 proteins, respectively.

**Video S7**:

Subcellular localization of 35S::GFP:NPH3-4K/A in hypocotyl cells of stably transformed Arabidopsis *nph3-7* upon BL-irradiation.

**Video S8**:

Subcellular localization of 35S::GFP:NPH3ΔC51 in hypocotyl cells of stably transformed Arabidopsis *nph3-7* upon BL-irradiation.

**Video S9**:

Subcellular localization of 35S::GFP:NPH3ΔC28 in hypocotyl cells of stably transformed Arabidopsis *nph3-7* upon BL-irradiation.

**Video S10**:

Dynamic BL-induced changes in the subcellular localization of 35S::GFP:NPH3ΔN54 in hypocotyl cells of stably transformed Arabidopsis *nph3-7*.

## References

1. Motchoulski, A., and Liscum, E. (1999). Arabidopsis NPH3: A NPH1 photoreceptor-interacting protein essential for phototropism. Science 286, 961–964.

2. Christie, J.M., and Murphy, A.S. (2013). Shoot phototropism in higher plants: new light through old concepts. Am J Bot 100, 35–46.

3. Fankhauser, C., and Christie, J.M. (2015). Plant phototropic growth. Current biology : CB 25, R384–389.

4. Legris, M., and Boccaccini, A. (2020). Stem phototropism toward blue and ultraviolet light. Physiol Plant 169, 357–368.

5. Inoue, S., Kinoshita, T., Matsumoto, M., Nakayama, K.I., Doi, M., and Shimazaki, K. (2008). Blue light-induced autophosphorylation of phototropin is a primary step for signaling. Proc Natl Acad Sci U S A 105, 5626–5631.

6. Mackintosh, C. (2004). Dynamic interactions between 14-3-3 proteins and phosphoproteins regulate diverse cellular processes. Biochem J 381, 329–342.

7. Sullivan, S., Kharshiing, E., Laird, J., Sakai, T., and Christie, J.M. (2019). Deetiolation Enhances Phototropism by Modulating NON-PHOTOTROPIC HYPOCOTYL3 Phosphorylation Status. Plant Physiol 180, 1119–1131.

8. Rakusová, H., Fendrych, M., and Friml, J. (2015). Intracellular trafficking and PIN-mediated cell polarity during tropic responses in plants. Curr Opin Plant Biol 23, 116–123.

9. Willige, B.C., Ahlers, S., Zourelidou, M., Barbosa, I.C., Demarsy, E., Trevisan, M., Davis, P.A., Roelfsema, M.R., Hangarter, R., Fankhauser, C., et al. (2013). D6PK AGCVIII kinases are required for auxin transport and phototropic hypocotyl bending in Arabidopsis. Plant Cell 25, 1674–1688.

10. Ding, Z., Galván-Ampudia, C.S., Demarsy, E., Łangowski, Ł., Kleine-Vehn, J., Fan, Y., Morita, M.T., Tasaka, M., Fankhauser, C., Offringa, R., et al. (2011). Light-mediated polarization of the PIN3 auxin transporter for the phototropic response in Arabidopsis. Nat Cell Biol 13, 447–452.

11. Zourelidou, M., Absmanner, B., Weller, B., Barbosa, I.C., Willige, B.C., Fastner, A., Streit, V., Port, S.A., Colcombet, J., de la Fuente van Bentem, S., et al. (2014). Auxin efflux by PIN-FORMED proteins is activated by two different protein kinases, D6 PROTEIN KINASE and PINOID. Elife 3.

12. Christie, J.M., Suetsugu, N., Sullivan, S., and Wada, M. (2018). Shining Light on the Function of NPH3/RPT2-Like Proteins in Phototropin Signaling. Plant Physiol 176, 1015–1024.

13. Hohm, T., Preuten, T., and Fankhauser, C. (2013). Phototropism: translating light into directional growth. Am J Bot 100, 47–59.

14. Pedmale, U.V., Celaya, R.B., and Liscum, E. (2010). Phototropism: mechanism and outcomes. Arabidopsis Book 8, e0125.

15. Inada, S., Ohgishi, M., Mayama, T., Okada, K., and Sakai, T. (2004). RPT2 is a signal transducer involved in phototropic response and stomatal opening by association with phototropin 1 in Arabidopsis thaliana. Plant Cell 16, 887–896.

16. de Carbonnel, M., Davis, P., Roelfsema, M.R., Inoue, S., Schepens, I., Lariguet, P., Geisler, M., Shimazaki, K., Hangarter, R., and Fankhauser, C. (2010). The Arabidopsis PHYTOCHROME KINASE SUBSTRATE2 protein is a phototropin signaling element that regulates leaf flattening and leaf positioning. Plant Physiol 152, 1391–1405.

17. Lariguet, P., Schepens, I., Hodgson, D., Pedmale, U.V., Trevisan, M., Kami, C., de Carbonnel, M., Alonso, J.M., Ecker, J.R., Liscum, E., et al. (2006). PHYTOCHROME KINASE SUBSTRATE 1 is a phototropin 1 binding protein required for phototropism. Proc Natl Acad Sci U S A 103, 10134–10139.

18. Pedmale, U.V., and Liscum, E. (2007). Regulation of phototropic signaling in Arabidopsis via phosphorylation state changes in the phototropin 1-interacting protein NPH3. J Biol Chem 282, 19992–20001.

19. Tsuchida-Mayama, T., Nakano, M., Uehara, Y., Sano, M., Fujisawa, N., Okada, K., and Sakai, T. (2008). Mapping of the phosphorylation sites on the phototropic signal transducer, NPH3. Plant Science 174, 626–633.

20. Haga, K., Tsuchida-Mayama, T., Yamada, M., and Sakai, T. (2015). Arabidopsis ROOT PHOTOTROPISM2 Contributes to the Adaptation to High-Intensity Light in Phototropic Responses. Plant Cell 27, 1098–1112.

21. Kimura, T., Haga, K., Nomura, Y., Higaki, T., Nakagami, H., and Sakai, T. (2020). The Phosphorylation Status of NPH3 Affects Photosensory Adaptation During the Phototropic Response. bioRxiv.

22. Jaspert, N., Throm, C., and Oecking, C. (2011). Arabidopsis 14-3-3 proteins: fascinating and less fascinating aspects. Frontiers in Plant Science 2, 96.

23. Keicher, J., Jaspert, N., Weckermann, K., Möller, C., Throm, C., Kintzi, A., and Oecking, C. (2017). Arabidopsis 14-3-3 epsilon members contribute to polarity of PIN auxin carrier and auxin transport-related development. Elife 6.

24. Bustos, D.M., and Iglesias, A.A. (2006). Intrinsic disorder is a key characteristic in partners that bind 14-3-3 proteins. Proteins 63, 35–42.

25. Piovesan, D., Necci, M., Escobedo, N., Monzon, A.M., Hatos, A., Mičetić, I., Quaglia, F., Paladin, L., Ramasamy, P., Dosztányi, Z., et al. (2021). MobiDB: intrinsically disordered proteins in 2021. Nucleic acids research 49, D361–d367.

26. Mergner, J., Frejno, M., List, M., Papacek, M., Chen, X., Chaudhary, A., Samaras, P., Richter, S., Shikata, H., Messerer, M., et al. (2020). Mass-spectrometry-based draft of the Arabidopsis proteome. Nature 579, 409–414.

27. Wang, P., Hsu, C.C., Du, Y., Zhu, P., Zhao, C., Fu, X., Zhang, C., Paez, J.S., Macho, A.P., Tao, W.A., et al. (2020). Mapping proteome-wide targets of protein kinases in plant stress responses. Proc Natl Acad Sci U S A 117, 3270–3280.

28. Coblitz, B., Wu, M., Shikano, S., and Li, M. (2006). C-terminal binding: an expanded repertoire and function of 14-3-3 proteins. FEBS letters 580, 1531–1535.

29. Johnson, C., Crowther, S., Stafford, M.J., Campbell, D.G., Toth, R., and MacKintosh, C. (2010). Bioinformatic and experimental survey of 14-3-3-binding sites. Biochem J 427, 69–78.

30. Kansup, J., Tsugama, D., Liu, S., and Takano, T. (2014). Arabidopsis G-protein beta subunit AGB1 interacts with NPH3 and is involved in phototropism. Biochem Biophys Res Commun 445, 54–57.

31. Heilmann, I. (2016). Phosphoinositide signaling in plant development. Development 143, 2044–2055.

32. Platre, M.P., Noack, L.C., Doumane, M., Bayle, V., Simon, M.L.A., Maneta-Peyret, L., Fouillen, L., Stanislas, T., Armengot, L., Pejchar, P., et al. (2018). A Combinatorial Lipid Code Shapes the Electrostatic Landscape of Plant Endomembranes. Dev Cell 45, 465–480 e411.

33. Simon, M.L., Platre, M.P., Marques-Bueno, M.M., Armengot, L., Stanislas, T., Bayle, V., Caillaud, M.C., and Jaillais, Y. (2016). A PtdIns(4)P-driven electrostatic field controls cell membrane identity and signalling in plants. Nat Plants 2, 16089.

34. Tejos, R., Sauer, M., Vanneste, S., Palacios-Gomez, M., Li, H., Heilmann, M., van Wijk, R., Vermeer, J.E., Heilmann, I., Munnik, T., et al. (2014). Bipolar Plasma Membrane Distribution of Phosphoinositides and Their Requirement for Auxin-Mediated Cell Polarity and Patterning in Arabidopsis. Plant Cell 26, 2114–2128.

35. Gronnier, J., Crowet, J.M., Habenstein, B., Nasir, M.N., Bayle, V., Hosy, E., Platre, M.P., Gouguet, P., Raffaele, S., Martinez, D., et al. (2017). Structural basis for plant plasma membrane protein dynamics and organization into functional nanodomains. Elife 6.

36. Inoue, S., Kinoshita, T., Takemiya, A., Doi, M., and Shimazaki, K. (2008). Leaf positioning of Arabidopsis in response to blue light. Mol Plant 1, 15–26.

37. Barbosa, I.C., Shikata, H., Zourelidou, M., Heilmann, M., Heilmann, I., and Schwechheimer, C. (2016). Phospholipid composition and a polybasic motif determine D6 PROTEIN KINASE polar association with the plasma membrane and tropic responses. Development 143, 4687–4700.

38. Brzeska, H., Guag, J., Remmert, K., Chacko, S., and Korn, E.D. (2010). An experimentally based computer search identifies unstructured membrane-binding sites in proteins: application to class I myosins, PAKS, and CARMIL. J Biol Chem 285, 5738–5747.

39. Perraki, A., Cacas, J.L., Crowet, J.M., Lins, L., Castroviejo, M., German-Retana, S., Mongrand, S., and Raffaele, S. (2012). Plasma membrane localization of Solanum tuberosum remorin from group 1, homolog 3 is mediated by conformational changes in a novel C-terminal anchor and required for the restriction of potato virus X movement]. Plant Physiol 160, 624–637.

40. Banani, S.F., Lee, H.O., Hyman, A.A., and Rosen, M.K. (2017). Biomolecular condensates: organizers of cellular biochemistry. Nat Rev Mol Cell Biol 18, 285–298.

41. Zavaliev, R., Mohan, R., Chen, T., and Dong, X. (2020). Formation of NPR1 Condensates Promotes Cell Survival during the Plant Immune Response. Cell 182, 1093–1108 e1018.

42. Kimura, T., Tsuchida-Mayama, T., Imai, H., Okajima, K., Ito, K., and Sakai, T. (2020). Arabidopsis ROOT PHOTOTROPISM2 Is a Light-Dependent Dynamic Modulator of Phototropin1. Plant Cell 32, 2004–2019.

43. Emenecker, R.J., Holehouse, A.S., and Strader, L.C. (2021). Biological Phase Separation and Biomolecular Condensates in Plants. Annual review of plant biology.

44. Ruff, K.M., Roberts, S., Chilkoti, A., and Pappu, R.V. (2018). Advances in Understanding Stimulus-Responsive Phase Behavior of Intrinsically Disordered Protein Polymers. Journal of molecular biology 430, 4619–4635.

45. Kaiserli, E., Sullivan, S., Jones, M.A., Feeney, K.A., and Christie, J.M. (2009). Domain swapping to assess the mechanistic basis of Arabidopsis phototropin 1 receptor kinase activation and endocytosis by blue light. Plant Cell 21, 3226–3244.

46. Sakamoto, K., and Briggs, W.R. (2002). Cellular and subcellular localization of phototropin 1. Plant Cell 14, 1723–1735.

47. Preuten, T., Blackwood, L., Christie, J.M., and Fankhauser, C. (2015). Lipid anchoring of Arabidopsis phototropin 1 to assess the functional significance of receptor internalization: should I stay or should I go? New Phytol 206, 1038–1050.

48. Cheng, Y., Dai, X., and Zhao, Y. (2007). Auxin synthesized by the YUCCA flavin monooxygenases is essential for embryogenesis and leaf formation in Arabidopsis. Plant Cell 19, 2430–2439.

49. Furutani, M., Kajiwara, T., Kato, T., Treml, B.S., Stockum, C., Torres-Ruiz, R.A., and Tasaka, M. (2007). The gene MACCHI-BOU 4/ENHANCER OF PINOID encodes a NPH3-like protein and reveals similarities between organogenesis and phototropism at the molecular level. Development 134, 3849–3859.

50. Treml, B.S., Winderl, S., Radykewicz, R., Herz, M., Schweizer, G., Hutzler, P., Glawischnig, E., and Ruiz, R.A. (2005). The gene ENHANCER OF PINOID controls cotyledon development in the Arabidopsis embryo. Development 132, 4063–4074.

51. Furutani, M., Nakano, Y., and Tasaka, M. (2014). MAB4-induced auxin sink generates local auxin gradients in Arabidopsis organ formation. Proc Natl Acad Sci U S A 111, 1198–1203.

52. Furutani, M., Sakamoto, N., Yoshida, S., Kajiwara, T., Robert, H.S., Friml, J., and Tasaka, M. (2011). Polar-localized NPH3-like proteins regulate polarity and endocytosis of PIN-FORMED auxin efflux carriers. Development 138, 2069–2078.

53. Glanc, M., Van Gelderen, K., Hoermayer, L., Tan, S., Naramoto, S., Zhang, X., Domjan, D., Včelařová, L., Hauschild, R., Johnson, A., et al. (2021). AGC kinases and MAB4/MEL proteins maintain PIN polarity by limiting lateral diffusion in plant cells. Current biology : CB.

54. Blatt, M.R., and Grefen, C. (2014). Applications of fluorescent marker proteins in plant cell biology. Methods Mol Biol 1062, 487–507.

55. Binder, A., Lambert, J., Morbitzer, R., Popp, C., Ott, T., Lahaye, T., and Parniske, M. (2014). A modular plasmid assembly kit for multigene expression, gene silencing and silencing rescue in plants. PLoS One 9, e88218.

56. Hecker, A., Wallmeroth, N., Peter, S., Blatt, M.R., Harter, K., and Grefen, C. (2015). Binary 2in1 Vectors Improve in Planta (Co)localization and Dynamic Protein Interaction Studies. Plant Physiol 168, 776–787.

57. Karimi, M., Depicker, A., and Hilson, P. (2007). Recombinational cloning with plant gateway vectors. Plant Physiol 145, 1144–1154.

58. Julkowska, M.M., Rankenberg, J.M., and Testerink, C. (2013). Liposome-binding assays to assess specificity and affinity of phospholipid-protein interactions. Methods Mol Biol 1009, 261–271.

59. Park, M., Touihri, S., Muller, I., Mayer, U., and Jurgens, G. (2012). Sec1/Munc18 protein stabilizes fusion-competent syntaxin for membrane fusion in Arabidopsis cytokinesis. Dev Cell 22, 989–1000.

60. Albert, I., Böhm, H., Albert, M., Feiler, C.E., Imkampe, J., Wallmeroth, N., Brancato, C., Raaymakers, T.M., Oome, S., Zhang, H., et al. (2015). An RLP23–SOBIR1–BAK1 complex mediates NLP-triggered immunity. Nature Plants 1, 15140.

61. Schindelin, J., Arganda-Carreras, I., Frise, E., Kaynig, V., Longair, M., Pietzsch, T., Preibisch, S., Rueden, C., Saalfeld, S., Schmid, B., et al. (2012). Fiji: an open-source platform for biological-image analysis. Nat Methods 9, 676–682.

62. Zavaliev, R., and Epel, B.L. (2015). Imaging callose at plasmodesmata using aniline blue: quantitative confocal microscopy. Methods Mol Biol 1217, 105–119.

